# Structure of HIV-1 Env glycoprotein on virions reveals an alternative fusion subunit organization and native membrane coupling

**DOI:** 10.64898/2026.01.09.698652

**Authors:** Jacob T. Croft, Hung N. Do, Daniel P. Leaman, Klaus N. Lovendahl, Pooja Ralli-Jain, Katelyn J. Chase, Chengbo Chen, Vidya Mangala Prasad, Cynthia A. Derdeyn, Michael B. Zwick, S. Gnanakaran, Kelly K. Lee

**Affiliations:** Department of Medicinal Chemistry, University of Washington, Seattle, WA, USA; Theoretical Biology and Biophysics Group, Theoretical Division, Los Alamos National Laboratory, Los Alamos, NM, USA 87545; Department of Immunology and Microbiology, Scripps Research, La Jolla, CA, USA; Department of Laboratory Medicine & Pathology, University of Washington, Seattle, Washington 98195, United States; Molecular Biophysics Unit, Indian Institute of Science, Bangalore, India

**Keywords:** Human Immunodeficiency Virus type-1, envelope protein, cryo-electron tomography, Hydrogen/Deuterium-exchange mass spectrometry, molecular dynamics, *in situ* structure, subtomogram averaging, membrane-proximal external region, membrane, virus, glycoprotein

## Abstract

An effective vaccine for Human Immunodeficiency Virus type-1 (HIV-1) has yet to be developed, and detailed characterization of functional Env glycoprotein, the primary antigenic target on virions, has remained elusive. While engineered Env trimers recapitulate many aspects of functional Env, key differences in antigenicity and dynamic behavior have been reported. Here, cryo-electron tomography and subtomogram averaging of HIV-1 virus-like particles (VLPs) revealed conformational differences in critical membrane-proximal regions compared to soluble Envs. Hydrogen/Deuterium-Exchange Mass Spectrometry and Molecular Dynamics captured dynamic profiles of membrane-bound Env and identified critical interactions with membrane. We show that disruption of the viral membrane results in relaxation of Env to a form that resembles engineered, soluble trimers. Additionally, Env from mature and immature VLPs exhibit only minor conformational differences, while surface clustering on virions changes significantly. These studies provide new insights into the essential role the membrane plays in maintaining Env in its native conformational form.

## Introduction

An effective HIV-1 vaccine that can prevent infection by inducing broadly neutralizing antibodies (bnAbs) against diverse primary isolates remains an important public health goal. Despite the availability of powerful anti-retroviral combination therapies, over one million new infections take place each year ^1^. The HIV-1 envelope glycoprotein (Env) has remained a challenging target for vaccine design due to the instability of its native trimeric state and extreme variation among circulating viral isolates ^2^. Moreover, recombinant Env tends to misfold, misassemble, and aggregate, resulting in antigens that are poor facsimiles of the native Env trimer, unless extensively engineered and modified to lock in specific conformations.

For decades, intensive efforts have focused on engineering the Env trimer to maintain the stable “prefusion” form as a prerequisite for eliciting HIV-1 bnAbs that could either block CD4 receptor/coreceptor engagement, and/or prevent conformational changes that drive membrane fusion and genome delivery ^3–8^. Indeed, the vast majority of structural insight into the Env has been based upon studies of soluble trimeric ectodomains engineered with stabilizing mutations that lock in what has been assumed to be the Env prefusion conformation, or Env that has been extracted from membranes and solubilized with detergent, bicelles, or reconstituted into nanodiscs that do not necessarily retain the natural lipid composition or leaflet asymmetry found in biological membranes ^5,9–12^. Biophysical studies have shown, however, that such trimers, while closely mimicking native Env ^5^, exhibit notable differences in dynamic properties and epitope exposure ^13,14^. Moreover, the membrane-interactive regions, transmembrane domain and large cytoplasmic tail are often truncated, even though these regions are known to modulate ectodomain conformation, antigenicity, and stability ^15^.

A comprehensive analysis of functional trimer on virions has remained elusive, though cryo-electron tomography (cryo-ET) approaches have started to resolve Env *in situ* ^16–18^. An accurate picture of the native antigenic target that elucidates the display of antibody epitopes as they exist on the virion surface can inform the design of improved vaccine immunogens that are more likely to elicit effective antibody responses. Notably, recent vaccine studies comparing soluble and membrane-bound forms of Env have shown that membrane presentation produced a quantitatively and qualitatively improved immune response ^19,20^. Additionally, determination of the structure of the prefusion form of Env, including its membrane interactive components, is critical to understand how it engages with the viral membrane and functions as a viral fusion protein that mediates the viral genome’s delivery into cells.

Studies have indicated that Env presentation on a biological membrane may stabilize a key antigenic form that changes when membrane interactions are disrupted by perturbing the membrane or by mutating membrane-interactive residues ^15,21^. This could be mediated in part by the membrane-proximal external region (MPER), a key motif in the gp41 subunit and highly conserved target for some of the most cross-reactive HIV-1 bnAbs identified to date ^22^. Indeed, numerous biochemical studies suggest the MPER and associated parts of Env can modulate its assembly, global behavior, and antigenicity ^15,21–25^. Membrane presentation also affects steric accessibility of membrane-proximal epitopes due to its ability to tilt ^12,17,26,27^. To date, however, it has been difficult to resolve membrane-interactive regions of Env.

Here, we used cryo-electron tomography with subtomogram averaging combined with molecular dynamics (MD) and hydrogen-deuterium exchange mass spectrometry (HDX-MS) to analyze Env on VLP from multiple HIV-1 isolates including those derived from transmitted/founder variants. From this integrative study, we constructed a structural model for fully functional HIV-1 Env trimers on the surface of mature HIV-1 virion. We resolved an unexpected organization for the gp41 fusion subunit that differs noticeably from the configuration that has been observed in nearly all Env trimer structures to date including those of full-length, non-engineered trimers solubilized with detergent or nanodiscs ^9,11,12,26,27^. Our analysis revealed the configuration of membrane interactions mediated by MPER segments as well as identifying a previously unrecognized interaction involving the conserved C-terminus of the gp120 subunit. Asymmetric behavior among protomers in the trimers and relation to the membrane interactions and trimer tilting were analyzed. Critically, we demonstrated that when membrane coupling was disrupted, Env ectodomain relaxes to the SOSIP-like trimer conformation. These studies demonstrate that presentation of Env on a native viral membrane determines the organization and conformational regulation of this important antigenic target.

## Results

### An alternative organization for membrane-interactive regions of gp41 in virus-bound Env

To determine the structure of Env *in situ*, virus-like particles (VLPs) were first produced using a cell line stably expressing high levels of a partial cytoplasmic tail truncated Env (ADA.CM.755*) ^28^, and transfection of an Env-deficient proviral plasmid backbone, as described previously ^17^. The VLPs produced from these cells incorporate high levels of Env that can be readily visualized by cryo-electron tomography (cryo-ET). We performed subtomogram averaging and resolved the structure of Env to 8.6 Å resolution with the application of C3 symmetry (Fig 1A, Fig S1). While much of the resulting density map was consistent with previously determined structures of closed, prefusion Env, displaying especially good agreement in the organization of the gp120 subunits (Fig 1A, Fig S2A-C), the gp41 subunit differed substantially relative to previously reported structures (Fig 1B, S2D) ^9,11,12,26,27^. Furthermore, the current structure offers notable improvement for ADA.CM.755* Env compared to a previous study ^17^, with higher resolution, more complete density for all structural elements including for gp41 and membrane-interactive regions.

**Figure 1.**
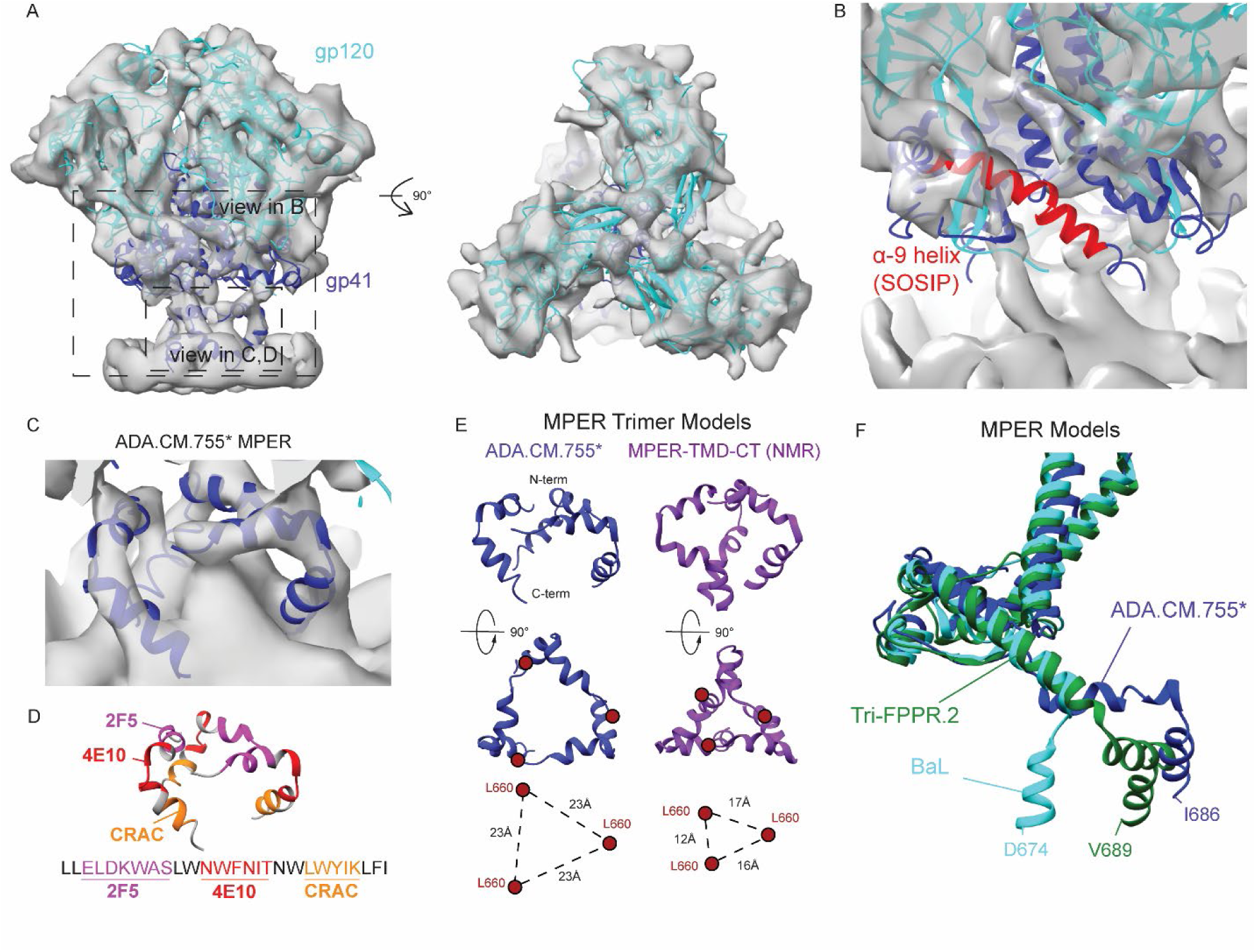
HIV-1 Env structure in the viral membrane. **(A)** Cryo-ET density map (transparent grey) of HIV-1 ADA.CM.755* Env determined by subtomogram averaging of Env on the viral membrane with gp120 (cyan) and gp41 (blue) models fitted to the density. View in panel B shown in box. **(B)** Close up on the structure of a soluble Env SOSIP construct (PDB: 4ZMJ) fit into the cryo-ET density with the gp41 HR2 α-9 helix colored. The SOSIP conformation of the α-9 helix does not fit the electron density of ADA.CM.755* Env on the virus surface. **(C)** Membrane-proximal density attributed to the MPER sequence, with a model of the MPER fitted in. The model was refined from the NMR structure of a MPER-TM-CT construct (PDB: 7LOI) ^30^. **(D)** MPER structure annotated with the 2F5 epitope (pink), 10E8 epitope (red) and LWYIK/CRAC CHOL-binding motif (orange). **(E)** Comparison of the trimeric MPER model fitted to the ADA.CM.755* cryo-ET density map (dark blue) to the trimeric MPER model determined by NMR of a MPER-TM-CT construct (purple, PDB: 7LOI) ^30^. **(F)** Comparison of single chain MPER conformations. For each model, a single chain of gp41 is shown. ADA.CM.755* Env is colored dark blue, BaL Env structure determined by subtomogram averaging is cyan ^16^, and Tri-FPPR.2 Env is colored green ^12^. The relative orientations of the MPER models were determined by aligning the ectodomain structures, except for the MPER-TM-CT NMR model which was manually aligned to the ADA.CM.755* density.

In the gp41 configuration first reported for native-like trimer ectodomains and solubilized Env trimers, the α-9 helix in the HR2 segment of gp41 that connects the ectodomain to MPER and membrane forms a long, extended helix ^29^. In our new reconstruction, density for an ordered helix was not clearly resolved (Fig 1B, S2D). Refinement with C1 symmetry resulted in differences in the density of this region between subunits, but the α-9 helix was still not clearly resolved in any of the subunits (Fig S1E-F). These data indicated that this segment of gp41 either exhibited significant conformational heterogeneity between particles or the polypeptide was in an extended or flexible conformation in VLP-associated Env.

While the α-9 helix density was missing, a substantial leg of density was observed extending into the membrane beneath each protomer. By constructing a pseudoatomic model with flexible fitting to our reconstructed map, we determined that the segment was consistent with a helix-turn-helix motif, corresponding to significant portions of the MPER segment, which fit neatly into the leg of density that descends into the membrane from beneath the *adjacent* protomer positioned counter-clockwise to the first protomer (viewed from above the membrane) (Fig 1C). This segment can then be connected to last portion of gp41 density by modeling a portion of the α-9 segment as a loop.

In this model, the N-terminal half of MPER containing the bnAb 2F5 epitope lay approximately parallel to and in association with the outer membrane leaflet, with the bnAb 10E8 epitope forming a bend in the helix, while the C-terminal segment containing a “LWYIK” Cholesterol (CHOL) binding motif (AKA cholesterol recognition amino acid consensus or “CRAC motif”) descended at an angle into the membrane (Fig 1D). Together, the three MPER subunits formed a set of struts beneath the Env ectodomain that is reminiscent of the “mace”-shaped trimeric MPER structure determined by NMR spectroscopy for a bicelle-solubilized, truncated MPER-TM-CT construct (Fig 1E) ^30^. In comparison to the NMR structure, in which the N-terminal L660 residues of MPER of the construct converge at the center of the trimer (separated by 12-17Å), in ADA.CM.755* Env this segment exhibited an outward rotation allowing for connectivity with the ectodomain (Fig 1E), with the L660 residues located 23 Å apart. This structure also differs from a recently reported asymmetric reconstruction of a stabilized “Tri-FPPR” Env construct reconstituted in A18 lipid nanodiscs ^12^, in which the MPER of one subunit was positioned more distally, directly connecting to a canonical SOSIP-like ordered α-9 helix (Fig 1F).

Notably, all of these MPER configurations contrasted with a structure determined by subtomogram averaging of Env from a neutralization-sensitive, lab-adapted HIV-1 strain, BaL ^16^. We built the MPER N-terminus into the BaL Env map, resulting in a model in which the α-9 helix takes on the SOSIP-like configuration, consistent with previous modeling ^16^, followed by a sharp turn at its terminus that positions the MPER N-terminal segment in a three-helix bundle directly perpendicular to the membrane (Fig 1F). In contrast to the other MPER models in which this structural element lies in close association with the membrane, the BaL MPER appears to sit largely atop the membrane, indicating a substantially different configuration and exposure relative to what we observe in ADA.CM.755* Env ^16^.

### HIV-1 variant-specific differences in gp41 configuration

Due to the differences between gp41 conformation in our ADA.CM.755* model and the SOSIP-like conformation in other published structures (Fig 1B, 2A, S2D), we sought to determine whether the lack of ordering of the α-9 helix was a peculiarity of ADA.CM Env or a result of the partial truncation of the cytoplasmic tail. To test this, we expressed several different fully functional Envs from a range of isolates on VLPs and generated subtomogram-averaged structures for comparison with known trimer structures and the ADA.CM.755* Env reconstruction (Fig 2B-K, Fig S3).

**Figure 2.**
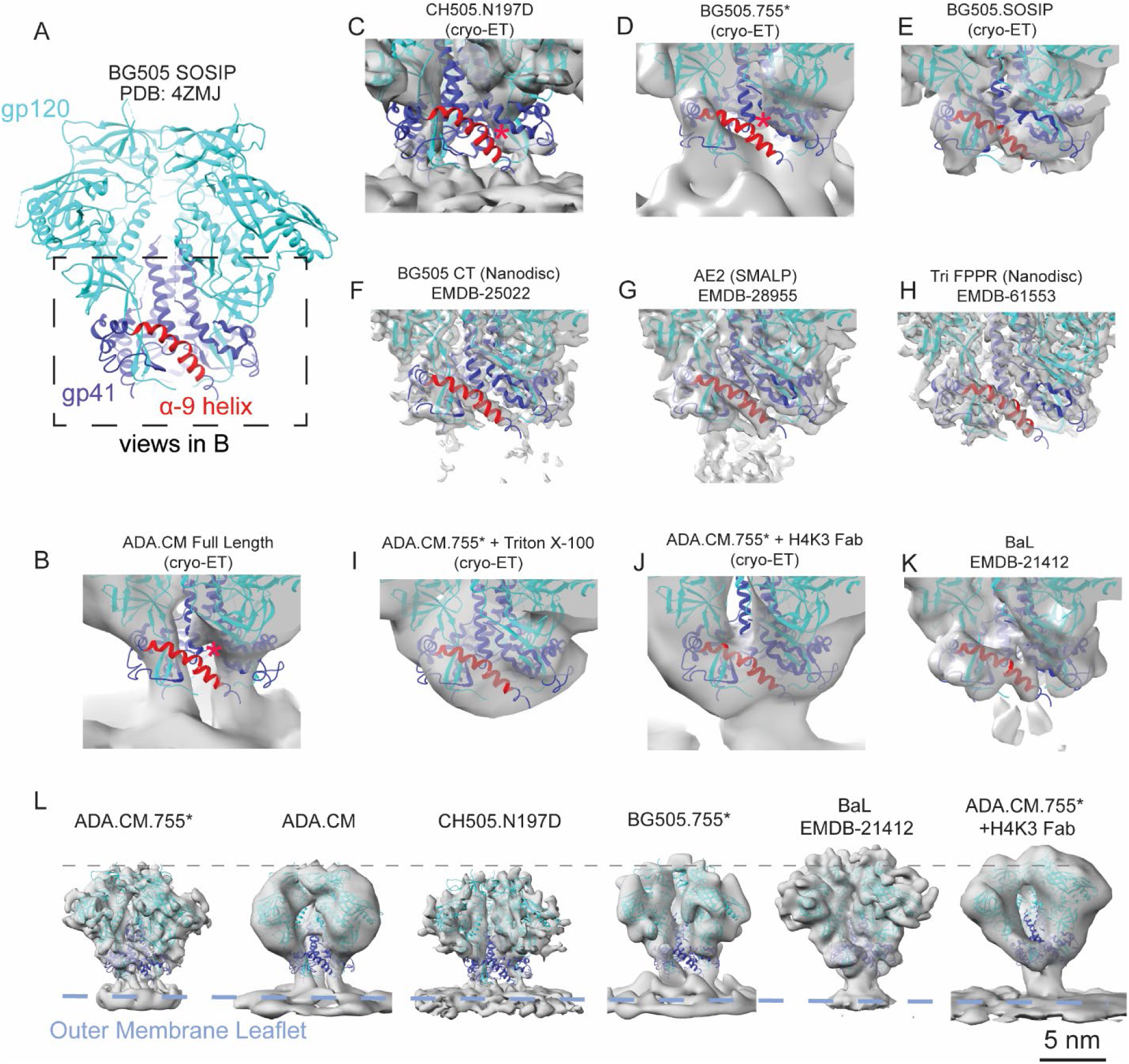
HR2 α-9 helix conformation is dependent on MPER embedding on membrane. **(A)** Model of BG505 SOSIP with gp120 colored cyan, gp41 colored dark blue, and the α-9 helix colored red. **(B-K)** Model from panel B fit into electron densities for a panel of Env determined in this study and others ^11,12,26,102^. Red asterisk indicates models in which the α-9 helix does not fit the density **(L)** Comparison of Env ectodomain maps determined by subtomogram averaging. Maps were aligned on the density of their outer membrane leaflet, and the difference in height to the trimer apex was determined by fitting a model of the ectodomain to each map. ADA.CM.755*, and ADA.CM, BG505.755* were all <1 Å different in height. ADA.CM.755* Env treated with MPER antibody PGZL1.H4K3 Env was slightly tilted so the height of each subunit differed. BaL Env is raised 8.5 Å from the membrane.

First, we examined full-length ADA.CM Env including the complete cytoplasmic tail. This comparison is of interest since the tail has been reported in certain circumstances to influence Env’s antigenic profile ^31^. Similar to the ADA.CM.755* Env (Fig 1B), the α-9 helix from the SOSIP configuration falls outside of the density (Fig 2B). We also investigated another full-length Env, CH505.N197D, which is incorporated on VLPs at higher levels than full-length ADA.CM, allowing the ectodomain structure to be solved to higher resolution (9.2 Å) (Fig 2C, S3). The CH505.197D Env density map was consistent with the same HR2 segment being flexible or conformationally variable and exhibited even less density for gp41 than the other constructs, indicating a high degree of conformational heterogeneity within the membrane-proximal portions of gp41 for this Env. Next, we determined the structure of BG505.755* Env to 17Å resolution. This Env bears the same ectodomain sequence as the BG505 SOSIP that was the first and most thoroughly characterized native-like Env trimer ^5,29,32,33^. However, in this case, the SOS disulfide bond and I559P substitution that are defining SOSIP modifications were not included, while the MPER, TMD, and CT up to residue 755 were included. Similar to ADA.CM.755* Env, BG505.755* exhibited a similar organization with a lack of clear density where the α-9 helix would normally be positioned as in the SOSIP configuration (Fig 2D).

We questioned whether the absence of density in these regions resulted from subtomogram averaging and resolution limitations, reconstruction procedure, or may be due to imaging conditions. To address this, we expressed soluble BG505.SOSIP.664 and performed subtomogram averaging on the native-like trimer ectodomain using the same approach used for the VLP-displayed Env. In this case, the density map showed much more substantial density that encloses the volume where the α-9 helix is positioned (Fig 2E). We therefore conclude that for the set of diverse, fully functional Envs on VLP that we examined the α-9 helix is differently configured than in SOSIP and solubilized trimers, and the resolution achieved by our subtomogram averaging approach is sufficient to resolve differences in this region.

### The native gp41 configuration is dependent on interaction with viral membrane

Since the α-9 helix is also resolved in single-particle reconstructions of Env in membrane-like environments such as micelles, nanodiscs, and styrene-maleic acid lipid nanoparticles ^9,11,12^ (Fig 2F-H), we next investigated whether perturbation of the viral membrane would return Env to a SOSIP-like conformation. Treatment of immature (darunavir-treated) ADA.CM.755* VLPs with Triton X-100 resulted in membrane-disrupted immature cores that retained a population of Env^34,35^. Subtomogram averaging of this detergent-treated Env was consistent with the SOSIP structure (Fig 2I). Additionally, we performed subtomogram averaging on Env from ADA.CM.755* VLPs incubated with PGZL1.H4K3 (H4K3) Fab, which binds the MPER segment. Consistent with previous structures of MPER antibodies bound to Env ^9,27^, the ectodomain was tilted and partially extracted from the membrane, and its density accommodated an ordered α-9 helix, consistent with a SOSIP-like configuration (Fig 2J).

Notably, in the previously reported subtomogram average structure of native BaL Env on virus, the ectodomain was observed to exhibit a SOSIP-like conformation (Fig 2K) ^16^. Interestingly, the trimer apex of BaL Env appears to sit higher above the membrane than unliganded ADA.CM, ADA.CM.755*, CH505.N197D, and BG505.755* by approximately 8.5 Å (Fig 2L), apparently due to its exposed MPER conformation lifting the base of the ectodomain well-above the membrane. Measurements were made by manual alignment of the outer membrane leaflet in maps, and rigid fitting of the ectodomain structure (PDB 4ZMJ) ^10^ into each map followed by measurement of the distance between the same residue at the trimer apex between fits. Furthermore, we observed the MPER-Fab bound ADA.CM.755* Env not only exhibits a SOSIP-like ectodomain configuration with a predominant tilt as described above, but, like BaL Env, in this case, the ectodomain was observed to be lifted above the membrane to a similar degree as BaL Env (Fig 2L). Taken together, these results imply that the mode of interaction of MPER with the viral membrane is allosterically coupled with α-9 helix conformation, which takes on a stable SOSIP-like state upon partial or complete extraction from the viral membrane.

### Molecular Dynamics simulations reveal a flexible Env trimer base mediated by MPER and conserved gp120 C-terminus

To obtain a more detailed analysis of which residues and structural motifs are involved in coupling the Env ectodomain to the membrane, we performed atomistic (AA) MD simulations on fully glycosylated ADA.CM.755* and full-length CH505 Env embedded in a HIV-1 membrane mimic, with lipid composition based upon experimental HIV lipidomic analysis ^36,37^. We performed both density-guided MD simulations in which the Env models were dynamically fitted to our experimental cryo-ET densities as well as unbiased, conventional MD simulations. Coarse-grained (CG) MD simulations were also carried out from AA MD simulations to capture the complex lipid dynamics surrounding the Env. As part of a companion study, we also performed simulations with and without the immature MA lattice ^35^. Accounting for different HIV strains, systems, conditions, and replicates, a total of 48 AA simulations (with system sizes in the range of 1-1.5 million atoms) of at least one µs each and 8 CG simulations of at least 10 µs each were carried out.

In atomistic MD simulations, the HR2 α-9 segment of gp41 exhibited considerable flexibility, unlike in SOSIP structures, which supports our subtomogram averaging reconstructions in which a α-9 helix was not clearly resolved. While a long middle segment (640-647) of the α-9 maintained helicity, the beginning (629-634) and ending (648-655) segments of the α-9 exchanged between helices and loops (Fig 3A, S4A, C, E). Also, a short middle segment (635-638) was predominantly unstructured. Similarly, we observed that MPER sub-segments were flexible and exhibited conformational variability, predominantly helices and loops in the simulations, consistent with previously published NMR structure ^30^ (Fig 3B, S4B, D, F) and a recent MD simulation that explored the orientation of MPER in a truncated Env model ^38^. Secondary structures of the HR2 and MPER segments were asymmetric between the three protomers in the HIV-1 Env during the timescales of MD simulations (Fig S4).

**Figure 3.**
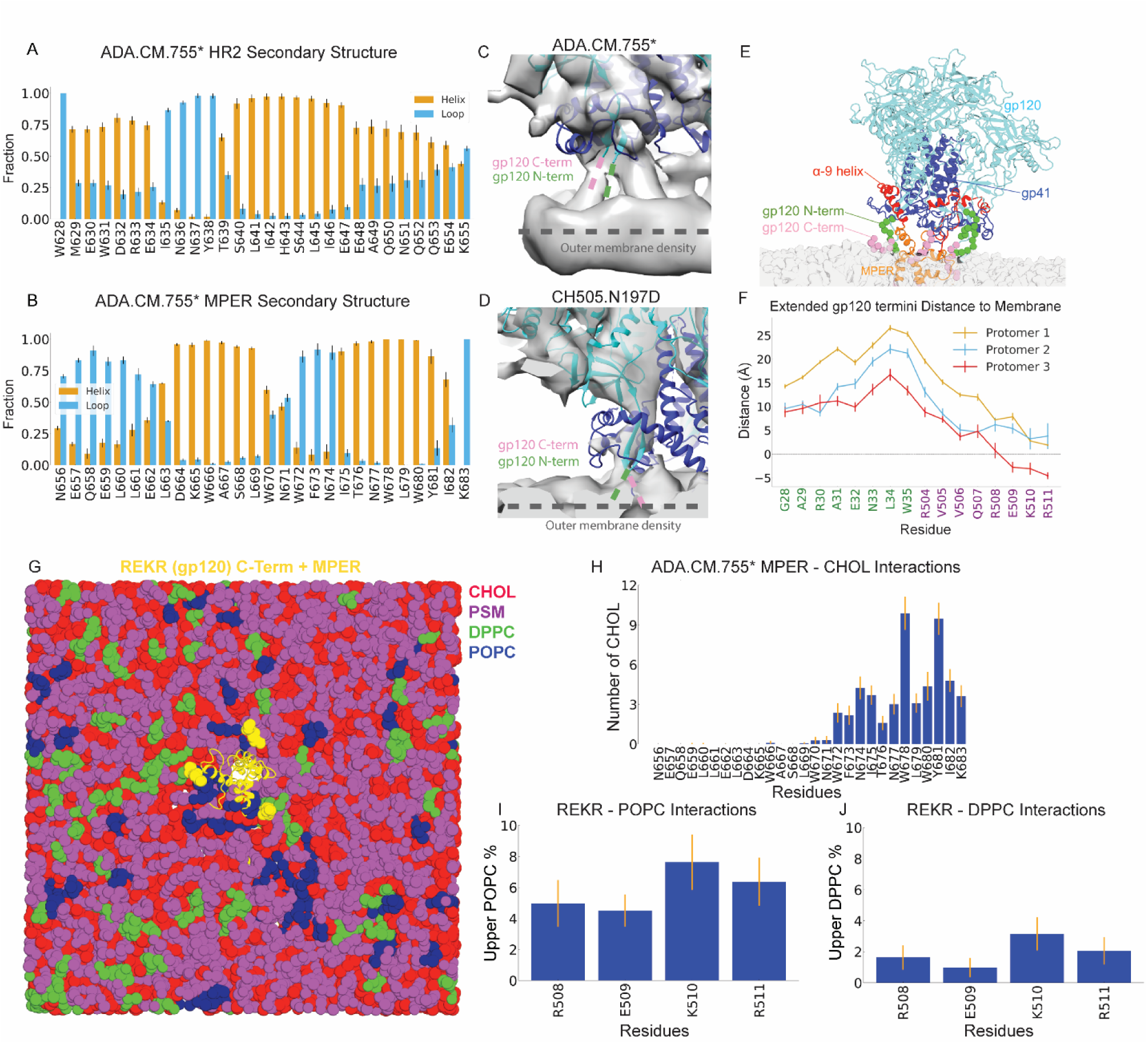
Molecular dynamics simulations of HIV-1 Env in modeled lipid bilayer. **(A-B)** Secondary structures of the **(A)** HR2 and **(B)** MPER residues sampled from the MD simulations (both density-guided and unbiased) of HIV-1 Env ADA.CM.755*. The fractions of simulation time the residues existed as helices are colored blue, while the fractions of simulation time the residues existed as loops are colored orange. **(C)** Close-up view on the ADA.CM.755* density map with the location of the gp120 N-terminus indicated by the dashed green line and the C-terminus indicated by the dashed pink line. **(D)** Close-up view on the CH505.N197D density map with the location of the gp120 N-terminus indicated by the dashed green line and the C-terminus indicated by the dashed pink line. **(E)** Insertion of the extended C-termini of gp120 of ADA.CM.755* into the membrane captured by structural modeling and MD simulations. The gp120 domain is colored cyan, gp41 ectodomain is colored blue, extended gp120 N-terminus is colored green, gp41 HR2 is colored red, MPER is colored orange, glycans are hidden, and the membrane is colored gray. **(F)** Average distances of the extended gp120 N- and C-termini relative to the phosphate lipid head groups in the membrane upper leaflet sampled from the MD simulations of ADA.CM.755* with extended gp120 termini in the model HIV-1 membrane. **(G)** Representative conformation of ADA.CM.755* with extended gp120 termini and MA in the HIV-1 membrane. Since the HIV-1 proteins were completely restrained during the CG simulations, the atomistic conformations of the MPER (shown as cartoon) and extended gp120 C-termini (shown as sphere) before the CG simulations were used for better visualizations. The HIV-1 proteins are colored yellow, CHOL lipid molecules are colored red, DPPC are colored green, POPC are colored blue, and PSM are colored magenta. **(H)** Interactions of the MPER residues of ADA.CM.755* with CHOL determined from the last 5µs of CG simulations of ADA.CM.755* in the HIV-1 membrane. **(I-J)** Enrichment of POPC **(I)** and DPPC **(J)** lipid molecules around the gp120 C-terminal residues (R508-R511) captured from the last 5µs of CG simulations of ADA.CM.755* in the HIV-1 membrane. A cutoff distance of 10 Å was used to determine the numbers and percentages of neighboring lipid molecules to each MPER residue. The error bars represent the standard errors of the means from different simulation replicas.

In addition to the membrane-interactive features attributed to the MPER in the ADA.CM.755* Env subtomogram averaged structure, additional shafts of density were observed spanning between the ectodomain and the viral membrane (Fig 3C). These are in close proximity to where the last-resolved residues in the C-terminal region of gp120 are positioned when available trimeric structures are docked into our density map. Similar connections between the ectodomain and the membrane, in close proximity to the gp120 termini were resolved in the subtomogram average of CH505.N197D Env, determined to 9.2 Å resolution (Fig 3D). We therefore hypothesized that the gp-120 C-terminus interacts directly with the membrane.

While the initial MD simulations did not include the gp120 C-termini, we examined the effect of extending the gp120 termini in our models past the residues that are usually resolved by cryo-EM and crystal structures and performed atomistic MD simulations of the membrane-embedded Env (Fig 3E, S5). We observed that the end of the extended C-terminus (RRVVQREKR) readily associated with the membrane, while the extended N-terminus (GARAENLWVTV) did not (Fig 3F, S5A). On average, residues G28-N33 of the extended gp120 N-termini were positioned away from the lipid head groups of the membrane upper leaflet, whereas residues R504-R511 of the extended gp120 C-termini were found proximal to the membrane. Association of the gp120 C-terminus to the membrane occurred in an asymmetric manner between subunits. Specifically, the basic residues K510 and R511 of protomer 3 were embedded in the upper leaflet of the membrane, possibly interacting with phosphate groups via electrostatic interactions (Fig. 3F, S5B). The interactions of the extended C-terminus with the membrane were more persistent than those of glycans. Consistent with a recent study ^39^, we observed the membrane proximity of two gp120 N-glycosylation sites (NGSs), N88 and N241, and four gp41 NGSs, N611, N616, N625, and N637 (Fig S5C-E). Within the limitations of the AA simulation timescales considered here, the observed glycan interactions with membranes, however, were found to be transient.

Next, to more comprehensively investigate lipid rearrangement around membrane-embedded Env, we carried out CG simulations of ADA.CM.755* and CH505 Envs, which allow for enhanced exploration of lipid mixing compared to AA MD simulations ^40–43^. The outer leaflet lipid distributions predominantly enriched in cholesterol (CHOL) and ordered lipids, palmitoyl sphingomyelin (PSM), and dipalmitoylphosphatidylcholine (DPPC), resembling a raft-like, ordered membrane composition showed lateral heterogeneity surrounding Env (Fig 3G, S6). In both ADA.CM.755* (Fig 3H) and CH505 (Fig S7), we observed that CHOL preferably interacted with L679, W680, Y681, I682, K683, i.e. the LWYIK motif ^44^, and a few additional residues preceding the motif. Indicative of a local ordered membrane environment, PSM interactions with MPER followed a similar profile as CHOL (Fig S6A-B, E-F). Interestingly, an enrichment of POPC (Fig 3I), a fluid-disordered lipid, over DPPC (Fig 3J) was seen around the region where the gp120 C-terminus made contact with the membrane, possibly enabling a more facile peripheral interaction with the basic residues.

### Maturation of the PR55^Gag^ lattice has a limited effect on Env structure and dynamics

Shortly after budding and release from the cell, HIV-1 particles undergo a proteolytic maturation step in which the internal PR55^Gag^ polyprotein is subjected to a series of cleavage events causing the lattice organization of the capsid and matrix proteins to rearrange ^45–47^. Previous reports have indicated that information about maturation state within the viral particle is transmitted through the viral membrane, likely via interaction of the Env cytoplasmic tail with matrix, resulting in differences in antigenicity of the gp41 subunit ^31^ as well as fusion activity of Env on immature HIV-1 particles ^48–50^. However, detailed structural analysis of Env on immature virus particles has not been reported to date.

We performed cryo-ET with subtomogram averaging on immature (darunavir-treated) VLPs bearing ADA.CM.755* Env and resolved the structure of the Env ectodomain to 9.7 Å resolution (Fig 4A, Fig S8). Similar to the mature structure (Fig 4B), immature Env exists in a closed, “hunkered-down” prefusion conformation with the SOSIP-like α-9 helix again absent in this case. While no major differences were observed between Gag maturation states, the maps exhibited slight differences in the connection of the ectodomain to the membrane. At low density threshold, the density attributed to the gp120 C-terminus connects to the membrane in both maps, but this feature and surrounding density is more robust in the immature map (Fig 4C-D). Additionally, density we attribute to MPER was not resolved as clearly in the immature map (Fig 4A,B).

**Figure 4.**
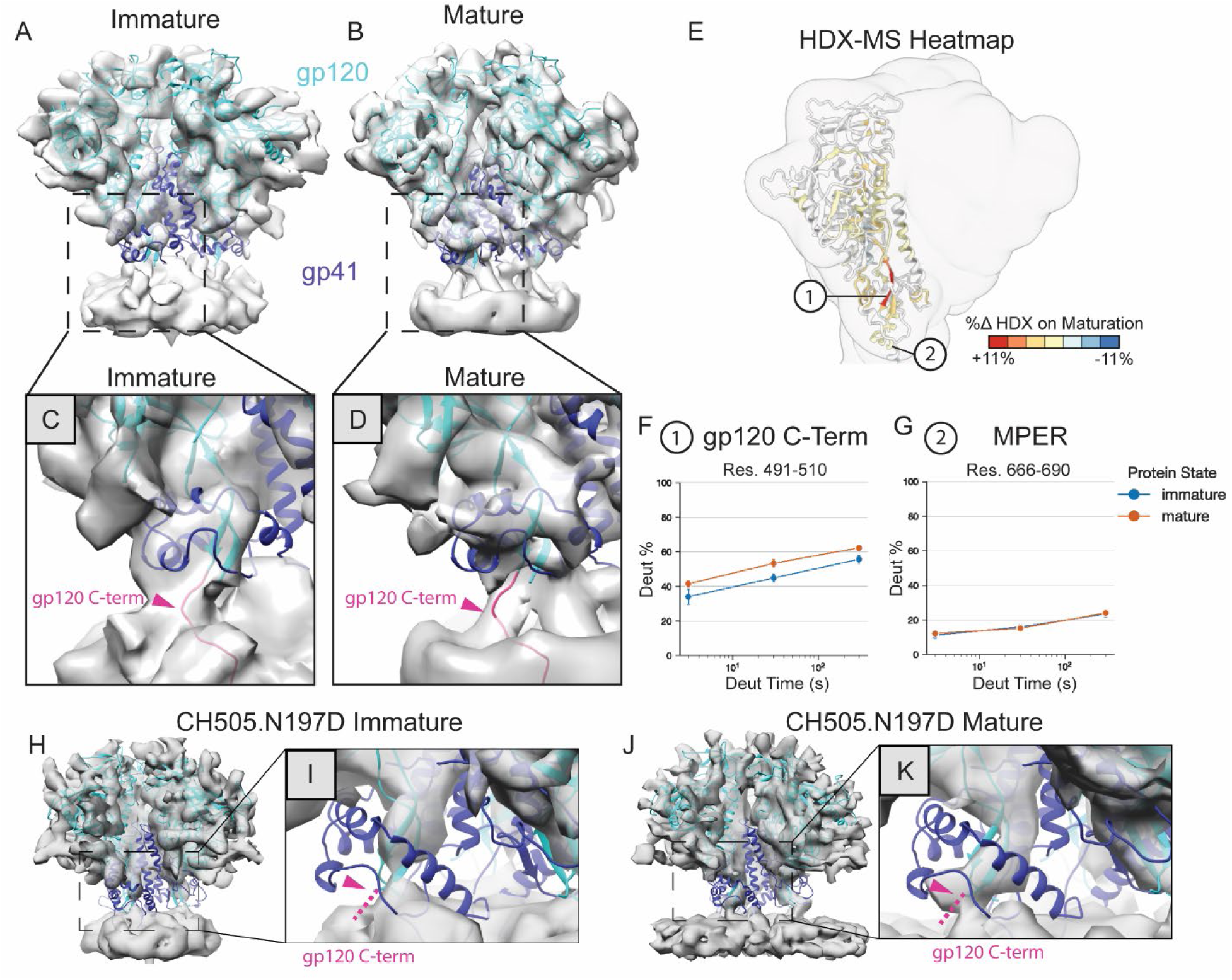
PR55^Gag^ maturation state has limited effect on Env ectodomain structure or dynamics. **(A, B)** Cryo-ET density maps (transparent grey) of HIV-1 Env ADA.CM.755* on immature **(A)** and mature (darunavir-treated) **(B)** virus-like particles determined by subtomogram averaging of Env on the viral membrane with gp120 (cyan) and gp41 (blue) model from the MD simulation of ADA.CM.755* Env with extended gp120 termini fitted to the density. **(C, D)** Close up view of gp41 and the gp120 termini-region. The extended gp120 C-terminal residues are colored magenta **(E)** Heatmap showing difference in protection by HDX-MS between ADA.CM.755* Env on mature and immature VLPs. Darker red regions are less protected in the mature sample, and darker blue regions are less protected in the immature sample. Grey regions indicate lack of coverage. Example peptides are indicated, in which uptake plots are shown for **(F)** the gp120 C-terminus which is more dynamic in mature VLPs and **(G)** MPER which does not show a differenc in uptake. Cryo-ET density maps (transparent grey) of HIV-1 Env CH505.N197D on immature **(H-I)** and mature (darunavir-treated) **(J-K)** virus-like particles determined by subtomogram averaging of Env on the viral membrane with the model from MD simulation of CH505 in complex with immature MA fitted to the density with gp120 colored cyan and gp41 colored blue. Panels **(I)** and **(K)** show views zoomed in on the gp120 termini and gp41 HR2 regions, with the location of the gp120 (residues not modeled) indicated with magenta arrowheads.

In order to test whether these subtle differences are real or a reflection of resolution limitations of the EM maps, HDX-MS was performed on both mature and immature ADA.CM.755* VLPs to assess local structural ordering and obtain a dynamic profile of the Envs *in situ*. This structural analytical method probes local protein backbone dynamic behavior and provides a sensitive fingerprint of a protein or assembly’s ordering and local flexibility. Consistent with our cryo-ET analysis, only minor differences were detected between the Env on the different particle types by HDX-MS (Fig 4E, S9). The only significant sites of different structural ordering localized to the gp120 C-terminus (Fig 4F). By HDX-MS, the gp120 C-terminus exhibited strong protection from solvent exchange in both cases, indicating involvement in 2° structure and/or sequestration from solvent. However, moderately greater protection from solvent exchange was detected for Env on immature particles. Furthermore, stronger cryo-ET density was resolved in this region in the immature VLP as well, consistent with greater ordering of the gp120 C-terminal membrane-interactive region. In contrast, we did not observe significant differences by HDX-MS for the MPER segment between maturation states (Fig 4G) which in both maturation states shows protection from exchange. This observation is consistent with the segment on average existing in an ordered helical configuration starting around residue 664, as would be in agreement with both the cryo-ET-based flexible fitting model (Fig 1D) and MD simulations (Fig 3B).

We also compared subtomogram averages of the Env ectodomain from immature (Fig 4H-I) and mature (Fig 4J-K) CH505.N197D VLPs, which include the complete cytoplasmic tail, to determine whether the lack of differences between maturation states was due to the CT truncation in ADA.CM.755* Env. At the resolution achieved (9.2 Å for mature, 8.9 Å for immature), no major differences were discernible in the maps. Similar to ADA.CM.755*, the gp120 termini were positioned near the outer membrane leaflet, which were connected in the density map.

### Env preferentially adopts a range of tilted conformations

In MD simulations, the Env ectodomain preferentially adopted conformations that were tilted with respect to the vertical z-axis of the viral membrane ^39^ (Figs 5A, S10A). First, we evaluated the differences in tilt-angle between immature (with MA lattice) and mature (without MA lattice) ADA.CM.755*. Though the tilt-angle distributions were similar between these two cases (Fig 5B, S10B), the distribution of mature ADA.CM.755* Env was slightly broader. However, the tilt-angle distribution of immature ADA.CM.755* Env, with and without the extended gp120 termini showed a noticeable difference (Fig 5B-C). We observed that the inclusion of the extended gp120 termini, resulted in a narrow distribution between 0° and 20°, with the most exhibiting tilts of 5°-10°, indicating that the association of the gp120 C-terminus with the membrane may have an influence on the Env ectodomain’s orientation.

**Figure 5.**
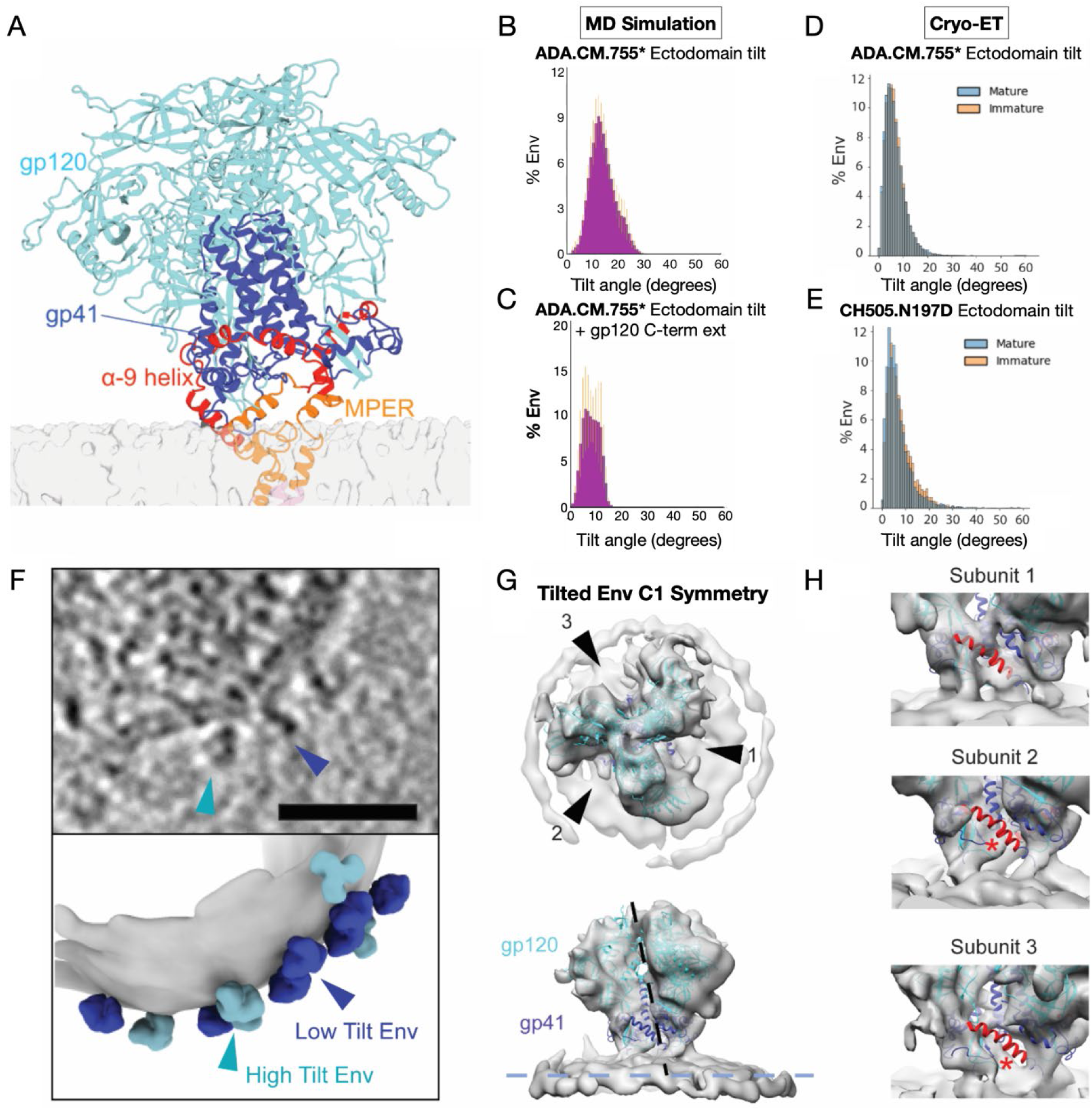
Tilting of the Env ectodomain in MD simulations and cryo-electron tomograms. **(A)** Representative conformation of the ectodomain of the HIV-1 Env ADA.CM.755* in complex with MA sampled from its density guided MD simulations. Subunit gp120 is colored cyan, subunit gp41 is colored cornflower blue, with its HR2 domain colored red, the MPER is colored orange, and the membrane is colored gray. **(B, C)** Distributions of the tilt-angles of the ectodomains relative to the z-axis (measured by the angle form by the vector drawn through the center-of-mass of the three helices formed by gp41 residues 570-592 and the vector drawn through the center of the simulation box) as shown in **Figure S10A** sampled from **(B)** the MD simulations of the glycosylated ADA.CM.755* envelope glycoprotein in complex with MA **(C)** and the MD simulations of the glycosylated ADA.CM.755* with extended gp120 termini in complex with MA. The error bars represent the standard error of the means from different simulation replicas. **(D-E)** Distribution of tilt-angles of particles by comparison of z axis orientation determined by focused refinement of the ectodomain vs the membrane during subtomogram averaging of Env on mature (blue) and immature (orange) VLPs of ADA.CM.755* Env **(D)** and CH505.N197D Env **(E)**. **(F)** Tomographic slice of an immature HIV-1 VLP bearing ADA.CM.755* Env (top). The bottom image shows a representation of the same VLP created by segmentation of the viral membrane (grey) and reprojection of both high-tilt (cyan) and low-tilt (blue) Env into the positions determined by subtomogram averaging. **(G)** High-tilt Env structure was determined by subtomogram averaging with C1 symmetry to 13.6 Å resolution. Views shown in panel G are indicated by arrowheads and the central axis of the ectodomain is shown by a dotted line. **(H)** Comparison of the density for each subunit in the reconstruction of tilted Env shown in panel F. Model of BG505 SOSIP (PDB: 4ZMJ) ^10^ is fit with gp120 colored cyan, gp41 colored dark blue, and the α-9 helix colored red. Red asterisks indicates subunits in which the α-9 helix does not fit the density. The subunit showing SOSIP-like α-9 helix density is located opposite the direction of Env tilting.

A wider range of ectodomain tilt-angles (0° to ∼55°) was observed for immature CH505 Env (Fig S10C). This result was consistent with the root-mean-square fluctuations (RMSF) calculations that indicated that CH505 Env exhibits greater flexibility relative to ADA.CM.755* (Fig S11). Unlike ADA.CM.755* Env, the tilt-angle distribution of mature CH505 was more restricted (0° to 40°) (Fig S10C-D) compared to the immature Env. We couldn’t detect a statistically significant correlation between glycan interactions with the membrane and Env tilting over the simulation timescales ^39^. Also, simulations of deglycosylated CH505 Env did not alter the range of sampled tilt-angles (Fig S10D-E). Extensive analysis of simulations (Fig 5A-C, S10) points towards the flexibility and conformational variability of Env regions at the solvent-membrane interface, such as α-9 helix and MPER, as possible drivers of Env ectodomain tilting.

We likewise compared the orientations of Env on individual VLPs in our subtomogram averages to the viral membrane and observed a similar distribution of tilt-angles from 0 to 25 degrees with a peak between 4 to 5 degrees (Fig 5D) with a similar distribution of tilting on immature particles (Fig 5D) and both maturation states of CH505.N197D (Fig 5E).

We classified Env on immature ADA.CM.755* VLPs (Fig S8) ^51^, identifying a class of Envs displaying higher tilt (Fig 5F) and obtained a tilted Env reconstruction at 13.6 Å resolution (Fig 5G, Fig S8). Notably, the tilted Env map contained differences in electron density between the three subunits near the connection to the membrane (Fig 5H). The subunit containing the α-9 helix directly opposite the direction of tilting towards the membrane (subunit 1) was most strongly resolved, while the α-9 helices of the other two subunits were not resolved. This is in accordance with our observation that membrane-extraction leads to stabilization of the α-9 helix in the SOSIP conformation, since the subunit with strong SOSIP-like α-9 helix density is pulled away from the membrane due to tilting. In contrast, tilting brings the other two subunits closer to the membrane, and they lack ordered α-9 helices. In agreement with these observations, in MD simulations, both the α-9 helix and MPER behaved asymmetrically among the three protomers. We thus conclude that tilting of Env is facilitated by structural plasticity of the gp41 α-9 and MPER regions.

## Discussion

A long-standing goal of HIV research has been to determine the structure of native Env on viral particles. Understanding the nature of the antigen in its native context can provide valuable insight to guide design of immunogens and to better understand mechanisms of neutralization by antibodies or inhibitors. Advances in engineering trimers for stability has led to hundreds of structures for the stabilized soluble ectodomains such as the widely studied SOSIP trimer design ^5^ and its derivatives. A number of structures have also been reported for recombinantly expressed Env extracted using detergents and nanodiscs ^9,11,12^, as well as an *in situ* structure of the lab-adapted strain BaL Env on mature virions ^16^. Remarkably, the reported structures exhibit excellent structural similarity between soluble and membrane bound forms with few exceptions. In contrast, here we observed differences in organization of the gp41 subunit of ADA.CM.755*, BG505.755* and full-length CH505.N197D Env on VLPs compared to previous structures. Collectively, we resolved density for the MPER region of ADA.CM.755* Env and observed apparent disordering of the α-9 segment across strains as well as apparent density connecting the gp120 C-terminus with the membrane. Our MD simulations supported these differences and revealed that the gp120 C-terminus, but not N-terminus, associates with the outer membrane leaflet.

Differences in Env conformational states have been indicated from single-molecule Förster resonance energy transfer (sm-FRET) experiments ^52–55^. sm-FRET revealed that native Env trimers on virus and SOSIP trimers sample at least three conformational states, distinguished by differences in FRET signals between two fluorescent reporters introduced onto specific sites on the trimer. Notably, in comparing matched BG505 SOSIP trimers vs BG505 Env on virions, the relative sub-populations of different FRET states as well as propensity to transition among the states were shown to be markedly different ^52,54^. It was proposed that Env on virions preferentially populates a conformational “state 1” that differs in significant ways from the SOSIP-like structure, which predominantly populates sm-FRET “state 2”. Most HIV-1 bnAbs by-and-large interact with both of these Env states, but some shift the population to state 1, while others bias Env to state 2 ^53,56,57^. A structural explanation for the observed differences has been missing, however, as all structures to date have yielded Env conformations that closely resemble the familiar SOSIP-type organization.

To explain why engineered trimers may differ in conformation and structural dynamic behavior from native, virus-displayed trimers, we note that the SOSIP trimers include a disulfide bond between gp120 and gp41 subunits, with one of the introduced cysteines, A501C, being located in gp120 C-terminal segment itself, which also is adjacent to HR2, one of the major sites showing differences between our models of Env on VLPs and previously determined structures. Likewise, the “IP” I559P substitution is situated in a loop region of helical region 1 (HR1) that likely re-directs its interactions with other parts of the trimer. Indeed, we see density for this loop reaching to contact the adjacent gp120 subunit in our reconstruction and models, which typically is not resolved in SOSIP structures.

Additionally, prior studies have drawn a link between Env antigenicity and properties of the viral membrane ^21,58^ which exhibits asymmetric outer and inner leaflet distribution and detergent-resistant, lipid raft-like compositions, including a significant enrichment in cholesterol ^37,59,60^. We conclude that differences between our structure and others result from a loss of native membrane coupling and specific Env-membrane interactions when Env is extracted from its native biological membrane environment. The interaction of MPER with lipid raft-like membrane enriched in CHOL appears to be a critical feature responsible for maintaining this coupling as suggested in previous reports and as our new intact Env in biologically realistic membrane simulations confirm ^44,61–63^. MPER mutations have also been shown to increase exposure of non-neutralizing epitopes which is partially reversed by “state 1” stabilizing mutations ^15^. In addition, when we perturb the MPER-membrane interaction via MPER antibody-binding or detergent extraction, the α-9 helix becomes ordered and takes on a helical SOSIP-like conformation (Fig 2I-J). Similarly, in the structure of highly tilted Env, we observed a similar ordering of the α-9 helix in the subunit that is pulled away from the membrane via tilting (Fig 5G). In accordance with this, the α-9 segment takes on a SOSIP-like conformation in the subtomogram average of the neutralization-sensitive BaL Env on virions, which displays an alternate MPER conformation in which the MPER N-terminus sits atop the membrane ^16^. Lastly, it is important to note that viral membrane-dependent conformational effects on Env are not fully recapitulated by detergent-solubilized or nanodisc-embedded Env, which also are seen to give rise to SOSIP-like conformations without clear resolution of MPER except in MPER-targeting Fab-bound conformations ^12,26,27^.

Several recent reports have suggested that Env preferentially adopts a tilted conformation ^11,12,17,27,39^. We also observed a range of Env tilting in our tomograms, as well as tilting of Env in MD simulations (Fig 5). The majority of Env particles were tilted to a less severe degree in our tomography data than the most frequent tilt-angle in MD simulations and the value reported by other reconstructions of nanodisc Env. Potentially, this could be due to an underestimation of tilting by our methods due to exclusion of higher-tilt Env during filtering of false positive particles in initial stages of subtomogram averaging. The lack of consensus and a well-defined coordinate for measuring the tilt-angle also adds ambiguity to reported values. Additionally, differences in membrane composition could affect the preferred tilt-angle of nanodisc-embedded vs native Env. Based upon our analysis, tilting is largely enabled by flexibility of the HR2 and MPER regions, which show different degrees of ordering among protomers when a given Env is tilted, as well as due to the engagement of the gp120 C-terminus, and, to a lesser degree, transient glycan interactions with the phospholipid headgroups. The observation that the gp120 C-terminus plays a role in modulating Env ectodomain coupling to the membrane has not been reported to date. This sequence is highly conserved presumably due to the role that furin cleavage plays, but it is conceivable that the role in grappling the ectodomain to the membrane and maintaining the conformation we observe may also impose selective pressure on this conserved region of the Env sequence. We also note that the gp120 C-terminal density along with membrane-proximal glycans appear to partially screen the conserved MPER epitope beneath the trimer.

In this study, we also report the first detailed analysis of Env on immature viral particles. Maturation of PR55^Gag^ within HIV-1 particles has been reported to affect Env gp41 antigenicity, with several epitopes including the C-terminal end of HR1, the fusion peptide proximal region (FPPR), and MPER being more exposed on immature HIV-1 particles ^31^. This suggested that Env may exist in an alternate conformation on immature viral particles. Additionally, immature HIV-1 particles fuse with receptor-bearing cells less efficiently than mature particles ^48–50^, leading to the hypothesis that Env may be held in an inactive conformation via the interaction of its cytoplasmic tail with the matrix domain of PR55^Gag^. Despite differences in interaction with MA between maturation states (described in accompanying paper)^35^, we only observed very minor differences in the ectodomain near the gp120 termini and connections to the membrane in ADA.CM.755* Env. This was corroborated by HDX-MS, which only detected a difference in protection at the gp120 C-terminus. We did not detect differences in MPER or the other regions previously shown to have altered antigenicity in immature particles ^31^. While lack of differences in ADA.CM.755* Env between maturation states could be due to CT truncation, we also did not resolve differences between the mature and immature CH505.N197D Env ectodomain. These subtle differences near the membrane are potentially due to differences in the organization of nearby lipids, which can be affected by the presence of the MA lattice ^35^. One possible explanation for reported differences in antigenicity is that Env retained on damaged, immature Gag lattices co-purified along with intact particles, which we frequently observe in immature virus samples, would lead to increased exposure of the MPER epitope and allow Env to relax to a distinct conformation compared to its membrane-bound form ^34^. Alternatively, stability of the specific Env being examined may be a factor; previous maturation studies examined a neutralization sensitive, lab-adapted isolate NL4-3, which is expected to be less stable than primary isolates and ADA.CM. 755* examined here ^64^. Lastly, as we discuss in the accompanying manuscript ^35^, Env distribution on the virus surface is highly dependent on maturation state, which could consequently influence avidity of antibodies, leading to differences in antigenicity.

Taken together, our study highlights differences in native, membrane-associated Env structure and dynamics compared to soluble or solubilized constructs. It also highlights the need to consider the effects of the viral membrane on Env conformation to enable successful presentation of the MPER epitope during vaccine design, as well as informing the development of more potent fusion inhibitors and use of MPER-targeting bnAbs as therapeutics. These studies indicate that the authentic membrane environment and native transmembrane and MPER sequences is critical for maintaining the native, prefusion conformation for the HIV-1 Env glycoprotein.

### Limitations of the Study

For biosafety reasons, mature ADA.CM.755*, CH505.N197D, and BG505 VLPs were deactivated with AT-2. After collection of mature ADA.CM.755* datasets, the 300 kV Titan Krios used for imaging was upgraded from a Gatan K2 to a Gatan K3 direct electron detector which was used for the immature ADA.CM.755*. This limits interpretation of subtle differences in quality of the maps. However, this should not affect the main finding that ectodomain displays a similar prefusion conformation in both maturation states. Additionally, the VLPs imaged by cryo-ET were produced in 293T cells, which may lead to a different membrane composition compared to virus in natural infection, and as is frequently done in pseudotyping HIV-1 particles, a mismatch of Env and Gag strains were present in the VLPs. Furthermore, we employed extensive multi-scale MD simulations to explore the ectodomain conformations of the pre-fusion HIV-1 Env glycoproteins and their interactions (including the glycans) with the HIV-1 membranes. Given the large size and complexity of membrane-bound HIV-1 Env systems, simulation timescales of 1 µs may be insufficient to capture their entire dynamics. In addition to force-field dependencies, there may be initial configuration biases. We established several protocols and measures to avoid those pitfalls. First, we performed multiple replicas of the simulations of the same system. Second, we conducted simulations of the same system with and without the experimental cryo-ET density constraints to assess any significant differences. Third, simulation-derived properties were reported with error bars. Finally, CG simulations with Martini 3 offer an excellent approach to sampling lipid mixing, with substantial simulation speedup due to its simplified representation, reduced degrees of freedom, and smoother energy landscape ^65^. We kept the protein component restrained during CG simulations due to limitations in accurately capturing the dynamics of large, flexible proteins with traditional Martini-based approaches. Ongoing efforts on that seek to integrate a virtual-site implementation of Go models with Martini 3 ^65^ should allow us to overcome this limitation in the future.

## Acknowledgements

This work was supported by National Institutes of Health grants R01-AI140868 (KKL) and R01-AI179697 (CAD, SG, KKL), R01-AI186650 (CAD, SG), Duke Center for HIV Structural Biology (U54-AI170752-01), and S10-OD032290. A portion of this work was also supported by grants from the Gates Foundation (INV-084290 and INV-010646) (KKL). We thank the University of Washington Arnold and Mabel Beckman Cryo-EM Center and the School of Pharmacy Mass Spectrometry Center for data collection time and support. We would also like to thank Max Crispin for providing unpublished data on CH505 Env glycosylation. Anju Yadav, Cesar Lopez and Mingfei Zhao for insights on setting up glycans and the complex membrane in the simulations. The authors acknowledge the support from the Los Alamos National Laboratory Institutional Computing for providing extensive computational resources.

## Author Contributions

Conceptualization, J.T.C., H.N.D., K.K.L. S.G.; Formal Analysis, J.T.C, H.N.D., K.N.L.; Funding Acquisition, K.K.L., S.G., C.A.D., M.B.Z.; Investigation, J.T.C., H.N.D., D.P.L., K.N.L., P.R.J., K.J.C., C.C., V.M.P.; Project Administration, K.K.L., S.G., C.A.D., M.B.Z.; Resources, M.B.Z; Software, J.T.C, H.N.D., K.N.L.; Supervision, K.K.L., S.G., C.A.D., M.B.Z.; Validation, J.T.C., H.N.D.; Visualization, J.T.C, H.N.D., K.N.L.; Writing, J.T.C., H.N.D., K.K.L. S.G.; Writing Review, C.A.D., M.B.Z., D.P.L., P.R.J., V.M.P., K.N.L.

## STAR Methods

### Key resources table

| REAGENT or RESOURCE | SOURCE | IDENTIFIER |
| --- | --- | --- |
| Chemicals, peptides, and recombinant proteins |  |  |
| Deuterium oxide 99.96% | Cambridge Isotope Laboratories | Cat #DLM-6-10X0.75 |
| 25 kDa Polyethylenimine (PEI) | Polysciences | Cat #23966 |
| Darunavir protease inhibitor | https://aidsreagent.org/ | ARP-11447 |
| Dulbecco's Modified Eagle Medium (DMEM) | ThermoFisher | Cat #10313-021 |
| Fetal Bovine Serum (FBS) | ThermoFisher | Cat #10437-028 |
| L-Glutamine | ThermoFisher | Cat #25030-081 |
| Penicillin Streptomycin | ThermoFisher | Cat #15140-122 |
| OptiPrep Density Gradient Medium (iodixanol) | Sigma | Cat #D1556 |
| Aldrithiol-2 (AT-2) | Sigma | Cat #143049 |
| HybridSPE-Phospholipid | Supelco | 55261-U |
| Nepenthesin-2 column, 2.1x20mm | AffiPro | AP-PC-004 |
| Deposited data |  |  |
| Subtomogram averaged EM density map: Mature ADA.CM.755* HIV-1 Env with C3 symmetry |  |  |
| Subtomogram averaged EM density map: HIV-1 ADA.CM.755* Env (Immature VLPs) |  | EMD-74813 |
| Subtomogram averaged EM density map: HIV-1 ADA.CM Env (Immature VLPs) |  | EMD-74789 |
| Subtomogram averaged EM density map: HIV-1 BG505.755* Env |  | EMD-74792 |
| Subtomogram averaged EM density map: HIV-1 ADA.CM.755* (Immature VLPs, Triton X-100 extracted) |  | EMD-74797 |
| Subtomogram averaged EM density map: HIV-1 Env BG505.SOSIP |  | EMD-74786 |
| Subtomogram averaged EM density map: HIV-1 ADA.CM.755* Env (Immature VLPs, tilted class) |  | EMD-74814 |
| Subtomogram averaged EM density map: HIV-1 CH505.N197D Env Ectodomain (Mature VLPs) |  | EMD-74763 |
| Subtomogram averaged EM density map: HIV-1 CH505.N197D Env Ectodomain (Immature VLPs) |  | EMD-74779 |
| PDB Deposition: Mature ADA.CM.755* HIV-1 Env by cryo-ET |  |  |
| PDB files from MD | Supporting Data 1 |  |
| Experimental models: Cell lines |  |  |
| ADA.CM.v4 (aka ADA.CM.755*.v4) in HEK239T stable cell line | 28 | N/A |
| ADA.CM.v2 (aka ADA.CM.755*.v2) in HEK239T stable cell line | 28 | N/A |
| CH505.N197D in HEK293T stable cell line | 66 | N/A |
| Recombinant DNA |  |  |
| pSG3ΔEnv Env-deficient HIV-1 backbone plasmid | https://aidsreagent.org/ | Cat#ARP-11051 |
| pIMC.CH505 | https://aidsreagent.org/ | Cat#ARP-13550 |
| Software and algorithms |  |  |
| SerialEM | 67 | https://bio3d.colorado.edu/SerialEM/ |
| Motioncor2 | 68 | https://emcore.ucsf.edu/ucsf-software |
| IMOD 4.11.15 | 69 | https://bio3d.colorado.edu/imod/ |
| EMAN2 (2.99.47) | 70 | https://blake.bcm.edu/emanwiki/EMAN2 |
| Topaz | 71 | https://emgweb.nysbc.org/topaz.html |
| UCSF Chimera | 72 | https://www.cgl.ucsf.edu/chimera/ |
| UCSF ChimeraX | 73 | https://www.rbvi.ucsf.edu/chimerax/ |
| Jupyter | 74 | https://jupyter.org/ |
| Byonic | Protein Metrics |  |
| HDExaminer | Sierra Analytics |  |
| SWISS-MODEL webserver | 75 | https://swissmodel.expasy.org/ |
| GROMACS 2024.5 | 76 | https://manual.gromgro.org/documentation/2024.5/index.html |
| AmberTools 2024 | 77 | https://ambermd.org/AmberTools.php |
| MDAnalysis | 78,79 | https://www.mdanalysis.org/ |
| PyMOL 3.1 | https://www.pymol.org/ | https://www.pymol.org/ |
| CHARMM-GUI webserver | 80–82 | https://charmm-gui.org/ |

## Experimental Models and Subject Details

### Cell lines for producing VLPs

Production of the “ADA.CM.v4” HEK293T cell line expressing high levels of HIV Env, was described previously ^28^. Cells were grown in D10 media (DMEM supplemented with 10% heat-inactivated fetal bovine serum (FBS; ThermoFisher Scientific), 20 mM L-glutamine, 100 U/mL penicillin, 100 mg/mL streptomycin, and 2.5 mg/mL puromycin (all media additives from ThermoFisher Scientific) and incubated at 37°C with 5% CO_2_). A stable cell line expressing Env CH505.N197D was generated by transduction of HEK293T cells using the lentiviral vector pLenti-III-HA following the same method as the ADA.CM.v4 cell line and has been described previously ^66^. The sex of the HEK293T cell line donor was female.

## Method Details

### VLP expression and purification

Stable cell lines expressing high levels of ADA.CM.755* and CH505.N197D Envs have been previously described ^28,66^. Cells were plated in 15-cm tissue culture dishes in 30 ml media (DMEM containing 10% heat-inactivated fetal bovine serum, 20 mM glutamine, 100 U/ml penicillin, 100 μg/ml streptomycin, and 2.5 μg/ml puromycin) at a cell density of 2.5 x 10^5^/ml and allowed to attach overnight. Next, 30 μg of Env-deficient HIV-1 backbone plasmid pSG3ΔEnv (NIH AIDS Reagent Program) in 900 μl DMEM was mixed with 90 μg 25 kDa PEI transfection reagent (Polysciences) in 900 μl DMEM, incubated for 15 minutes at room temperature, and the PEI/DNA mixture was added to the cells. Immature VLPs were produced by adding the protease inhibitor darunavir (NIH AIDS Reagent Program) to transfected cells at the same time as the DNA and PEI to a final concentration of 2 µM. Supernatants were harvested 3 days after transfection and cleared by centrifugation at 3,000 × *g* for 15 min. VLPs were pelleted at 40,000 × *g* for 1 h and resuspended 50-fold concentrated in PBS. VLPs were separated from cellular debris using iodixanol density gradient centrifugation. Concentrated VLPs were overlaid on a 9.6 to 20.4% iodixanol (Optiprep; Sigma) gradient, formed by layering iodixanol in 1.2% increments, and centrifuged at 200,000 × *g* for 1.5 h at 4°C in an SW41Ti rotor (Beckman). The top 4 ml of the gradient were discarded and then the next 5 ml were collected and pooled. Purified VLPs were brought up to 15 ml with PBS and concentrated to ∼0.2 ml using a 100 kDa MWCO Amicon centrifugal filter (Millipore) spun at 2,000 × *g*. VLPs were inactivated by adding aldrithiol-2 (AT-2) to a final concentration of 2.5 mM and samples were incubated at RT for 2 h. The volume was again increased to 15 ml with PBS and VLPs were concentrated using a 100 kDa MWCO centrifugal filter to 250-fold the concentration in the original transfection supernatant.

### Atomistic and coarse-grained simulations of envelope glycoproteins in the HIV-1 membranes

The results in this study were obtained from the atomistic (AA) simulations of the glycosylated ADA.CM.755* Env in complex with immature HIV-1 matrix (MA) protein, glycosylated ADA.CM.755* Env with extended gp120 termini in complex with immature HIV-1 MA protein, mature (without MA) glycosylated ADA.CM.755* Env, glycosylated CH505 Env in complex with immature HIV-1 MA protein, mature (without MA) glycosylated CH505 Env, and mature (without MA) deglycosylated CH505 Env as well as the coarse-grained (CG) simulations of the mature (without MA) CH505 Env and ADA.CM.755* Env with extended gp120 terminals in complex with immature HIV-1 MA protein embedded in the HIV-1 membrane lipid bilayers performed for a parallel study ^35^. The simulation protocols were summarized below.

The SWISS-MODELLER homology modeling webserver ^75^ was employed to model the full structures of the ADA.CM.755* Env and CH505 Env starting from their amino acid sequences. The ADA.CM.755* Env with extended gp120 termini were prepared by adding the GARAEN amino acid sequence before the LWVTV amino acid sequence of gp120 N-termini and adding the RRVVQREKR amino acid sequence after the VATKAK sequence of the gp120 C-terminus. The glycosylation profiles of the gp120 and gp41 ectodomains of the ADA.CM.755* Env were modeled based on the experimental glycosylation profile of the BG505 SOSIP.664 Env trimer ^83^.

The glycosylation profiles of the CH505 Env were modeled based on the unpublished experimental glycosylation profile of the CH505, along with the published profiles of BG505 SOSIP.664 Env trimer (cite 81). The structure of the immature HIV-1 matrix (MA) protein was taken from the 7OVQ PDB structure ^47^, with the central trimer and its three neighboring trimers used for simulations. The palmitoylation motif (CYSP) was attached to residues C764 and C843 in the gp41 cytoplasmic tail (CT) of CH505 Env, and the myristoylation motif (GLYM) was attached to residue G2 in each of the MA monomers in the immature MA lattice. The residues were numbered in accordance with the HXB2 numbering scheme of gp160 (https://www.hiv.lanl.gov/content/sequence/LOCATE/locate.html).

The AA simulation systems of ADA.CM.755* and CH505 Envs in complex with immature HIV-1 MA protein were prepared using the CHARMM-GUI webserver ^80–82^. The Envs were embedded in an asymmetric membrane composition mimicking the membrane lipid composition of HIV-1 virion ^36,37^. The outer leaflet was composed of 48% cholesterol (CHOL), 38% palmitoyl sphingomyelin (PSM), 9% dipalmitoylphosphatidylcholine (DPPC), and 5% palmitoyl-oleoyl-phosphatidylcholine (POPC). The inner leaflet was composed of 42% CHOL, 3% DPPC, 2% POPC, 4% palmitoyl-arachidonoyl phosphatidylethanolamine (PAPE), 4% palmitoyl-oleoyl phosphatidylethanolamine (POPE), 14% dioleoylphosphatidylethanolamine (DOPEE), 14% dipalmitoylphosphatidylethanolamine (DPPEE), and 17% palmitoyl-oleoyl-phosphatidylserine (POPS). The total number of lipid molecules in the upper leaflet was 864 molecules, while the total number of lipid molecules in the lower leaflet was 776 molecules in the HIV-1 membrane. Overall, the system sizes of the simulation systems ranged from ∼850,000 atoms (for mature Envs), with the cubic box dimensions of 207.5 Å × 207.5 Å × 207.5 Å, to ∼1 million atoms (for immature Envs with MA), with the box dimensions of 207.5 Å × 207.5 Å × 240.0 Å. The CHARMM36m force field parameter set ^84^ was used for the AA simulations.

Pulling simulations were carried out to insert the myristoylation motifs (GLYM) of the immature MA proteins, which were initially placed 20 Å from the gp41 CT, into the inner leaflet of the model HIV-1 membranes ^35^. The complexes of Envs and immature MA proteins, with the myristoylation motifs inserted in the membrane, were solvated in 0.15 M NaCl using the GROMACS 2024.5 ^76^ simulation package. The GROMACS 2024.5 ^76^ simulation package was employed to carry out the subsequent simulations of Envs in complex with immature MA ^35^. Energetic minimization, equilibrations with the constant number, volume, and temperature (NVT) and then with the constant number, pressure, and temperature ensembles, and a short 25-ns conventional MD (cMD) simulation was performed on all simulation systems, using timesteps of 1 fs for the NVT equilibration and 2 fs the NPT equilibration and cMD simulation. Four 100-ns density-guided combined with later unbiased production MD simulations and four sole unbiased production MD simulations were performed on the simulation systems of glycosylated ADA.CM.755* and CH505 Envs in complex with immature MA protein (in both cases of restraint and unrestraint MA), and four sole unbiased production MD simulations were performed on the other simulation systems, including mature (without MA) glycosylated ADA.CM.755* Env, glycosylated ADA.CM.755* Env with extended gp120 termini in complex with immature MA protein, mature (without MA) glycosylated CH505 Env, and mature (without MA) deglycosylated CH505 Env for up to 1µs per simulation replica, using a timestep of 2fs ^35^. The LINCS algorithm ^85^ was used to restrain all bonds containing hydrogen atoms and applied the periodic boundary conditions to our simulation systems. Furthermore, the velocity rescaling algorithm (cite 98 from the maturation paper) was employed to keep our simulation temperature constant at 310 K with a friction coefficient of 1.0 ps^-1^, and the stochastic cell rescaling algorithm (cite 99 from the maturation paper) with semi-isotropic coupling to keep pressure constant at 1.0 bar. Lastly, the pressure coupling constant was set to 5 ps, and the compressibility was set to 4.5 × 10^-5^ bar^-1^. For the density-guided simulations, the cross-correlation setting was used for the similarity measures, with atomic contribution to the experimental density set to be proportional to their masses. Furthermore, the density-guided force constant was set to 10^6^. The Gaussian transform spreading width was set to 0.2 nm, and the Gaussian transform spreading range in multiples of width was set to four. Our densities were set to be normalized and the adaptive force scaling was used, with an adaptive force scaling time constant set to 50 ps, to get much gentler fit in our density-guided simulations ^76^.

The CG simulations of mature (without MA) CH505 and ADA.CM.755* Envs with extended gp120 termini in complex with immature MA protein in model HIV-1 membranes, with four simulation replicas for each system, were set up to explore the lipid dynamics around the ectodomains of Env, starting from the final AA conformations ^35^. For the AA protein and membrane components of mature (without MA) CH505 and ADA.CM.755* with extended gp120 termini in complex with immature MA protein in model HIV-1 membranes, the models were converted to their coarse grained representations ^88^ using a combination of martinize2 ^89^ for the protein components, with modifications to accommodate for the myristoylation and palmitoylation motifs in the mapping files ^90^, and backward ^91^ python script for the lipid components. No glycans were backmapped due to the absence of their force field parameters in MARTINI 3 ^40^. The CG protein components were built with the side chain corrections ^92^ and elastic network ^93^. In particular, the elastic bond force constant was set to 1500 kJ.mol^-1^.nm^-2^ (-ef 1500), with the lower and upper bound of elastic bond cutoff set to 0 and 1.2 nm (−el 0 −eu 1.2). Meanwhile, the bond decay factor and power were set to 0 and 1.0 (−ea 0 −ep 1). The CG protein and membrane were solvated in 0.15 M NaCl solution using insane, with the box dimensions kept identical to the AA simulations ^94^. The MARTINI 3 force field parameter sets ^40^ were employed for the CG simulations to explore the lipid dynamics. The CG simulations were performed using the GROMACS 2024.5^76^ simulation software suite.

The CG simulation systems were also subjected to energetic minimization, followed by equilibration with NVT and then NPT ensembles, and one short 100-ns cMD simulation for equilibration ^35^. Four independent unbiased production MD simulations were performed on each simulation system for up to 10µs for each simulation replica, using a timestep of 10 fs. During the equilibration stage, the timestep was gradually increased from 2 fs during the equilibration with NVT ensemble to 5 fs and then 10 fs during the equilibration with NPT ensemble and cMD simulation. During the CG simulations, the protein components were completely restraint using a force constant of 250 kJ.mol^-1^.nm^-2^. The electrostatic interactions were calculated using the reaction field method ^95^ and Verlet cutoff scheme ^96^ with a cutoff distance of 1.1 nm for long-range interactions. Meanwhile, the temperature was kept constant at 310 K using the velocity rescaling thermostat ^86^ with a friction coefficient of 1.0 ps^-1^. Lastly, the pressure was kept constant at 1.0 bar using the stochastic cell rescaling ^87^ with semi-isotropic coupling, with the pressure coupling constant set to 5 ps and compressibility set to 3×10^-4^ bar^-1^.

At the end of the AA and CG simulations, the GROMACS 2024.5 ^76^ simulation software suite was employed to process the simulation trajectories, and the AmberTools ^77^, MDAnalysis ^78,79^ packages were used to analyze the simulation trajectories. In particular, MDAnalysis ^78,79^ was used in the calculations of secondary structures of HR2 and MPER, gp120 termini membrane insertion, glycan-membrane interactions, and ectodomain tilt-angles during the AA simulations as well as lipid distributions around the MPER and gp120 termini during the CG simulations. For the AA simulations, only the heavy atoms of the gp120 termini and glycans were used in calculating the distance between the gp120 residues as well as glycans and the phosphate lipid headgroups in the outer membrane leaflet. Furthermore, the tilt-angles of the ectodomains were calculated by measuring the angles formed by the vector drawn through the center-of-mass (COM) of the gp41 residues 570-592 (blue), the outer membrane center, and the z-axis. For the CG simulations, a contact definition of ≤ 10Å between any protein and lipid beads was used to calculate the lipid distributions around the MPER and gp120 termini. Meanwhile, AmberTools ^77^ was used in the calculations of Env RMSF during the AA simulations and hierarchical agglomerative clustering of the AA simulation frames to select representative AA conformations. In particular, hierarchical agglomerative clustering calculations were carried out to obtain the most populated conformations (whose fractions were ≥ 0.15) from the simulations of HIV-1 Env glycoproteins to serve as representative conformations. The clustering was carried out using the Cα-atom RMSD of HIV-1 Env gp120, gp41 ectodomain (including the HR2), and MPER residues relative to the starting conformations prior to the simulations as the distance metrics to cluster the HIV-1 Env conformations sampled in the imaged trajectories.

### Cryo-electron Tomography Data Collection

Tilt series of immature ADA.CM.755* VLPs, and ADA.CM VLPs were collected on a 300 kV Titan Krios G3 TEM (ThermoFisher) equipped with a K3 direct electron detector (Gatan) and GIF energy filter (Gatan) with slit width 20 eV at a magnification of 64,000x in super-resolution mode with an unbinned pixel size of 0.6932 e-/Å^2^. BG505 SOSIP, ADA.CM.755* VLPs treated with PGZL1.H4K3 FAb, and ADA.CM.755* VLPs treated with Triton X-100 were imaged on a 200 kV Glacios TEM (ThermoFisher) equipped with a K3 direct electron detector (Gatan) at 22,000x magnification in super-resolution mode with a pizel size of 0.93 e-/Å^2^ or in counting mode with a pixel size of 1.86 e-/Å^2^. BG505.755* VLPs on the same Glacios TEM with a K2 direct electron detector (Gatan) at 28,000x magnification and a pixel size of 1.4975 e-/Å^2^ in counting mode. Tilt series of mature ADA.CM.755* VLPs were collected as previously described ^17^ on a 300 kV Titan Krios G3 TEM (ThermoFisher) equipped with a K2 direct electron detector (Gatan) and GIF energy filter (Gatan) with slit width 20 eV. Unless otherwise specified tilt series were collected from either +/- 48 degrees or +/- 60 degrees with a step of 3 degrees with a dose symmetric tilt scheme using serialEM software ^67^ and a total dose of between 90-110 e-/Å^2^. Mature ADA.CM.755* VLPs were imaged with a total dose of 64-68 e-/Å^2^.

### Sample preparation for cryo-ET

All samples were prepared for cryo-ET by plunge freezing on a Vitrobot Mark IV (FEI). Three μL of purified VLPs were added to glow-discharged 300 mesh QF 1.2/1.3 or QF 2/2 grids (EMS and Ted Pella), blotted for 3-4 seconds with a blot force of zero, and plunge frozen in liquid ethane. For some samples, double blotting was used to increase concentration on grids. For these samples, 3 μL sample was applied, back-blotted away manually, 3 μL sample applied again, and then automatically blotted and plunge frozen in the vitrobot. The vitrobot was maintained at 4 C and 100% humidity during plunge freezing. Samples were either mixed with 10 nm BSA gold tracer at a ratio of 4:1 or 9:1, or frozen without gold beads. For experiments with PGZL1.H4K3 Fab, VLPs were treated with 8 μM Fab for 30 minutes prior to plunge freezing. For imaging of detergent-stripped VLPs, Triton X-100 was mixed with VLPs to a final concentration of 0.1%. After 15 min incubation on ice, the sample was run over a HiPPR detergent removal spin column (Thermofisher) and plunge frozen.

### Preprocessing and Tomogram Reconstruction

Cryo-ET data processing was performed as described in a parallel study ^35^. Prior to tomogram reconstruction, movies were motion corrected using MotionCor2 ^68^ and combined into ordered tilt-series stacks using the IMOD ^69^ newstack command. From this point onward, all subsequent processing was performed in EMAN2 (version 2.99.47) unless otherwise specified ^97^. Tilt series stacks were imported into EMAN2, the tomograms were aligned and reconstructed, and CTF estimation was performed. Some tomograms were denoised using Topaz prior to particle picking^71^. For all samples, particles were picked manually either in EMAN2 boxer or in IMOD slicer and imported into EMAN2.

### Mature ADA.CM.755* Subtomogram Averaging

Subtomogram averaging of ADA.CM.755* Env ectodomain is depicted in Fig S1. Preprocessing, tomogram reconstruction, and initial model generation was performed as described previously ^17^. The same particle set used in our previous study was reprocessed from this step onwards in EMAN2 ^97^ version 2.99.47 for comparison with the darunavir-treated ADA.CM.755* data processed in the same software, which lead to minor improvements in reported resolution and large improvement in map quality. First, the initial model was lowpass filtered to 50 Å and an initial refinement was performed with C1 symmetry and automasking, with resolution limited to 25 Å. All refinements in this study were performed using the “new” version of the refinement algorithm in EMAN2. This refinement was then used as an input for classification using EMAN2’s Gaussian mixture model (GMM) ^51^ with 5 classes. These classes were reconstructed without alignment and projected into the context of the tomograms for manual inspection. One class was discarded due to a noisy, low quality reconstruction, and another was discarded due to a higher percentage of erroneous particle orientation with respect to the viral particles. The remaining 3 classes were combined (19,083 particles) and both C1 and C3 symmetry refinements was performed limiting max resolution to 25 Å. Particles were then re-extracted, discarding duplicate particles and low cross-correlation particles (18,619 particles). Final, C1 and C3 symmetry refinements were performed on these particles using a resolution limit of 10 Å for alignment using a threshold mask containing just the ectodomain resulting in final map resolutions of 8.6 Å for both maps, reported using FSC=0.143 cutoff. Decreasing the resolution limit further did not lead to increase in resolution or map quality. Additionally, a C1-relaxed symmetry local refinement was performed starting from the parameters determined by the C3 refinement resulting in a final resolution of 9.4 Å, using FSC=0.143 cutoff.

### Immature (darunavir) ADA.CM.755* Subtomogram Averaging

Subtomogram averaging of ADA.CM.755* Env ectodomain from immature particles is depicted in Fig S8. After preprocessing, particles were extracted at 4X binning and an initial refinement was performed with C3 symmetry, with a resolution limit of 25 Å. Then, a local refinement was performed with C1 symmetry and the output of this was input into GMM classification ^51^ with 6 classes. Three classes (tilted and untilted Env) were selected based on particle orientation and map quality and another refinement was performed with C3 symmetry and a 10 Å resolution limit. Particles were re-extracted at binning of 2 and a local final refinement was performed with tight masking of the ectodomain resulting in a final resolution of 9.7 Å reported using FSC=0.143 cutoff.

A single class from the GMM classification was selected for processing of the tilted Env. C1 symmetry refinement was performed with a cylindrical mask encompassing the ectodomain and membranes. A tight mask over the ectodomain was created and final local refinement was performed of the tilted ectodomain resulting in a final resolution of 13.6 Å reported using FSC=0.143 cutoff.

### Miscellaneous Subtomogram Averaging

For BG505.755*, 4,826 particles were extracted at 4X binning and an initial refinement was performed with C3 symmetry, with a resolution limit of 25 Å using automasking. 4,496 particles were re-extracted at 2X binning and local refinement was performed with C3 symmetry, a resolution limit of 25 Å and using automasking. A final C3 refinement was performed using a tight mask over the ectodomain. A final resolution of 16.7 Å was reported using FSC=0.143 cutoff (Fig S3A).

For BG505.SOSIP, 50,329 particles were selected using EMAN2 convnet-based autopicking and extracted at 4X binning. An initial refinement was performed with C3 symmetry, with a resolution limit of 20 Å using automasking. Next 37,681 particles were extracted at 2X binning, and local refinement was performed with C3 symmetry. 2X-binned particles were once again filtered to a final set of 28.262 particles, and a final local refinement was performed resulting in an 8.0 Å map reported using FSC=0.143 cutoff (Fig S3B).

For full-length ADA.CM, 150 particles were extracted at 4X binning and global refinement was performed with C3 symmetry and a cylindrical mask resulting in a 25.1 Å map reported using FSC=0.143 cutoff (Fig S3C).

For Triton X-100 treated ADA.CM.755* VLPs, 809 Env particles were extracted at 4X binning and global refinement was performed with C3 symmetry and a cylindrical mask resulting in a 23.6 Å map reported using FSC=0.143 cutoff (Fig S3D).

For ADA.CM.755* + PGZL1.H4K3 FAb, 14,266 particles were extracted at 4X binning. The “new” initial model generation program was used in EMAN2 to generate an initial reference, which was low pass filtered to 50 Å. Global refinement was performed with C1 symmetry and a cylindrical mask resulting in 18.0 ang res map using FSC=0.143 cutoff (Fig S3E).

### Molecular modeling of ADA.CM.755* into moderate resolution cryo-ET density map

Homology models of the ADA.CM Env Ectodomain and MPER/TM/CT regions were made separately using SWISS-MODEL^75^ using PDB:8FAE (ectodomain) ^11^ and PDB:7LOI (MPER/TM/CT) ^30^ as templates. The ectodomain model was rigid-fit into the C3 symmetry cryo-ET density map using UCSF ChimeraX ^73^ and fit to the density using interactive molecular dynamics flexible fitting with ISOLDE ^98^. Due to the low resolution of the cryo-ET density, heavy distance restraints were applied to the model prior to flexible fitting. The homology model of the MPER region was manually fit to the cryo-ET density, and refined in ISOLDE restricting to the secondary structure present in the initial model. Finally, the two models were combined into a single model of the ectodomain and MPER regions.

### BG505.SOSIP Expression and Purification

The BG505 plasmid DNA construct (0.6 mg) and a furin expressing plasmid (0.2 mg) were co-transfected into 800 mL confluent Expi293F cell culture (3 x 10^6^ cells/mL) with ExpiFectamine 293 reagent following the ExpiFectamine 293 user guide. On the fifth day post-transfection, culture supernatant was collected from 30-minute centrifugation at 2000xg, followed by vacuum-filtration through 0.45 mm aPES filters. The filtered supernatant was supplemented with Tris-HCl (pH 7.4), EDTA and NaCl to match the composition of agarose bound Galanthus nivalis lectin (GNL) beads binding buffer (20 mM Tris-HCl, 1 mM EDTA, 120 mM NaCl, 0.02% NaN3, pH 7.4). For a 900-1000 mL supernatant, 5 mL GNL beads (50% slurry, VectorLabs) was added and incubated overnight at 4 °C. The GNL beads were then washed with 10 column volumes (CV) of binding buffer twice, followed by 2 CV of GNL elution buffer (1 M methyl α-D-mannopyranoside, 20 mM Tris-HCl, 1 mM EDTA, 120 mM NaCl, 0.02% NaN3, pH 7.4). Elution fractions were combined, buffer-exchanged into DEAE low-salt buffer (20 mM Tris-HCl, 100 mM NaCl, pH 7.4) and concentrated to ∼1.5 mg/mL with an Amicon Ultra-15-mL-centrifugal filter (100 K, Millipore). Concentrated sample was loaded onto pre-equilibrated 5-mL HiTrap DEAE column (Cytiva). Protein sample was collected from the flow-through of 20 mL low-salt buffer and the bound aggregates were later removed by high-salt buffer (20 mM Tris-HCl, 1 M NaCl, pH 7.4). Collected flow-through was buffer-exchanged to hydrophobic interaction column (HIC) binding buffer (2 M ammonium sulfate, 100 mM phosphate, pH 7.0) and loaded onto a pre-equilibrated 5-mL HiTrap Phenyl HIC column (Cytica). Step elution was applied with each 20 min elution of 1.5 M, 1 M, 0.5 M and 0 M ammonium sulfate containing buffer, respectively. Peak fractions from 1.5 M and 1 M ammonium sulfate elution, indicating BG505 trimers were combined, buffer-exchanged to HEPES buffer (10 mM HEPES, 200 mM NaCl, 0.02% NaN3, pH 7.5), and stored for experimental use.

### Env Tilting Analysis

For each cryo-ET dataset analyzed, after initial refinement of Env particles with a cylindrical mask, local refinements were performed masking the ectodomain and membrane individually. For the membrane refinements, translation of the particles was restricted to maximum 20 Å. The tilt-angle of the central axis of Env vs the normal to the membrane was calculated for each particle by computing the difference in particle orientation between the corresponding particle in both refinements.

### Hydrogen/deuterium-exchange Mass Spectrometry (HDX-MS)

HDX-MS experiments on the VLPs were performed as described previously with modifications ^17^. Purified mature & immature ADA.CM.V4 particles containing approximately 0.5mg/mL Env were desalted into HEPES-Buffered Saline (HBS, 10mM HEPES-NaOH, 150mM NaCl, pH 7.4) and the internal exchange standard PPPF added ^99^. Fifteen µL of VLP solution was diluted with 85µL of deuterated HBS and incubated at room temperature for the indicated time. The deuteration buffer contained angiotensin II and bradykinin peptides (AnaSpec) to serve as fully deuterated standards for measuring back-exchange.

Exchange was stopped by mixing with 100µL of chilled quench buffer (0.2M glycine-HCl, 4M guanidine-HCl, 0.2% DDM, 0.2M TCEP, pH 2.3) to bring the pH to 2.5. The mixture was incubated on ice for 30 sec before adding 20µL of ZrO2 beads (150mg/mL in 0.1M glycine-HCl), then incubated on ice for 1 min. The mixture was then transferred to a 0.45µm spin filter and centrifuged for 30 sec at 15,000 xg, 1°C. The final solution was transferred to a glass vial and frozen in a dry ice & ethanol bath, then stored at −80°C until analysis.

Samples were thawed and injected using a customized autosampler ^100^. Proteins were digested using a Nepenthesin-2 column kept at 15°C and separated on a C18 column using a 15 min gradient of acetonitrile in 0.1% formic acid. The protease column was cleaned after each injection using solutions of (1) Fos-choline-12 (Anaspec) in 0.1% TFA; (2) 2M GuHCl in 0.1% TFA; (3) 20% acetic acid, 5% ACN and 5% IPA (https://doi.org/10.1007/s13361-017-1860-3; https://doi.org/10.1007/s13361-012-0485-9). The trap column was washed after each injection to minimize sample carryover ^101^.

Peptides were analyzed using an Orbitrap Ascend mass spectrometer. Peptide identification from MS/MS data was performed using Byonic. Deuterium uptake analysis was performed with HD-Examiner. Data was visualized using HD-Examiner, Python, and ChimeraX.

### Quantification and Statistical Analysis

Representative images were chosen from cryo-ET analysis for figures.

## References

1. Global HIV & AIDS Statistics — 2025 Fact Sheet.

2. Barouch, D. H. Challenges in the development of an HIV-1 vaccine. Nature 455, 613–619 (2008).

3. Binley, J. M. et al. A recombinant human immunodeficiency virus type 1 envelope glycoprotein complex stabilized by an intermolecular disulfide bond between the gp120 and gp41 subunits is an antigenic mimic of the trimeric virion-associated structure. J. Virol. 74, 627–643 (2000).

4. Sanders, R. W. et al. Stabilization of the soluble, cleaved, trimeric form of the envelope glycoprotein complex of human immunodeficiency virus type 1. J. Virol. 76, 8875–8889 (2002).

5. Sanders, R. W. et al. A next-generation cleaved, soluble HIV-1 Env trimer, BG505 SOSIP.664 gp140, expresses multiple epitopes for broadly neutralizing but not non-neutralizing antibodies. PLoS Pathog. 9, e1003618 (2013).

6. Sharma, S. K. et al. Cleavage-independent HIV-1 Env trimers engineered as soluble native spike mimetics for vaccine design. Cell Rep. 11, 539–550 (2015).

7. Yang, L. et al. Structure-Guided Redesign Improves NFL HIV Env Trimer Integrity and Identifies an Inter-Protomer Disulfide Permitting Post-Expression Cleavage. Front. Immunol. 9, 1631 (2018).

8. Torrents de la Peña, A. & Sanders, R. W. Stabilizing HIV-1 envelope glycoprotein trimers to induce neutralizing antibodies. Retrovirology 15, 63 (2018).

9. Lee, J. H., Ozorowski, G. & Ward, A. B. Cryo-EM structure of a native, fully glycosylated, cleaved HIV-1 envelope trimer. Science 351, 1043–1048 (2016).

10. Kwon, Y. D. et al. Crystal structure, conformational fixation and entry-related interactions of mature ligand-free HIV-1 Env. Nat. Struct. Mol. Biol. 22, 522–531 (2015).

11. Wang, K. et al. Asymmetric conformations of cleaved HIV-1 envelope glycoprotein trimers in styrene-maleic acid lipid nanoparticles. Commun. Biol. 6, 535 (2023).

12. Qi, Y. et al. The membrane-proximal external region of human immunodeficiency virus (HIV-1) envelope glycoprotein trimers in A18-lipid nanodiscs. Commun. Biol. 8, 442 (2025).

13. Alsahafi, N. et al. SOSIP Changes Affect Human Immunodeficiency Virus Type 1 Envelope Glycoprotein Conformation and CD4 Engagement. J. Virol. 92, (2018).

14. Castillo-Menendez, L. R., Nguyen, H. T. & Sodroski, J. Conformational Differences between Functional Human Immunodeficiency Virus Envelope Glycoprotein Trimers and Stabilized Soluble Trimers. J. Virol. 93, (2019).

15. Wang, Q. et al. Global Increases in Human Immunodeficiency Virus Neutralization Sensitivity Due to Alterations in the Membrane-Proximal External Region of the Envelope Glycoprotein Can Be Minimized by Distant State 1-Stabilizing Changes. J. Virol. 96, e0187821 (2022).

16. Li, Z. et al. Subnanometer structures of HIV-1 envelope trimers on aldrithiol-2-inactivated virus particles. Nat. Struct. Mol. Biol. 27, 726–734 (2020).

17. Mangala Prasad, V., et al. Cryo-ET of Env on intact HIV virions reveals structural variation and positioning on the Gag lattice. Cell 185, 641–653.e17 (2022).

18. Li, W. et al. HIV-1 Env trimers asymmetrically engage CD4 receptors in membranes. Nature 623, 1026–1033 (2023).

19. Parks, K. R. et al. Vaccination with mRNA-encoded membrane-anchored HIV envelope trimers elicited tier 2 neutralizing antibodies in a phase 1 clinical trial. Sci. Transl. Med. 17, eady6831 (2025).

20. Ramezani-Rad, P. et al. Vaccination with mRNA-encoded membrane-bound HIV Envelope trimer induces neutralizing antibodies in animal models. Preprint at 10.1101/2025.01.24.634423 (2025).

21. Salimi, H. et al. The lipid membrane of HIV-1 stabilizes the viral envelope glycoproteins and modulates their sensitivity to antibody neutralization. J. Biol. Chem. 295, 348–362 (2020).

22. López, C. A., Alam, S. M., Derdeyn, C. A., Haynes, B. F. & Gnanakaran, S. Influence of membrane on the antigen presentation of the HIV-1 envelope membrane proximal external region (MPER). Curr. Opin. Struct. Biol. 88, 102897 (2024).

23. Nguyen, H. T. et al. Evaluation of the contribution of the transmembrane region to the ectodomain conformation of the human immunodeficiency virus (HIV-1) envelope glycoprotein. Virol. J. 14, 33 (2017).

24. Bradley, T. et al. Amino Acid Changes in the HIV-1 gp41 Membrane Proximal Region Control Virus Neutralization Sensitivity. EBioMedicine 12, 196–207 (2016).

25. Herschhorn, A. et al. The β20-β21 of gp120 is a regulatory switch for HIV-1 Env conformational transitions. Nat. Commun. 8, 1049 (2017).

26. Yang, S. et al. Dynamic HIV-1 spike motion creates vulnerability for its membrane-bound tripod to antibody attack. Nat. Commun. 13, 6393 (2022).

27. Rantalainen, K. et al. HIV-1 Envelope and MPER Antibody Structures in Lipid Assemblies. Cell Rep. 31, 107583 (2020).

28. Stano, A. et al. Dense Array of Spikes on HIV-1 Virion Particles. J. Virol. 91, (2017).

29. Pancera, M. et al. Structure and immune recognition of trimeric pre-fusion HIV-1 Env. Nature 514, 455–461 (2014).

30. Piai, A. et al. NMR Model of the Entire Membrane-Interacting Region of the HIV-1 Fusion Protein and Its Perturbation of Membrane Morphology. J. Am. Chem. Soc. 143, 6609–6615 (2021).

31. Joyner, A. S., Willis, J. R., Crowe, J. E., Jr. & Aiken, C. Maturation-Induced Cloaking of Neutralization Epitopes on HIV-1 Particles. PLOS Pathog. 7, 1–9 (2011).

32. Julien, J.-P. et al. Crystal structure of a soluble cleaved HIV-1 envelope trimer. Science 342, 1477–1483 (2013).

33. Lyumkis, D. et al. Cryo-EM structure of a fully glycosylated soluble cleaved HIV-1 envelope trimer. Science 342, 1484–1490 (2013).

34. Wyma, D. J., Kotov, A. & Aiken, C. Evidence for a Stable Interaction of gp41 with Pr55^Gag^ in Immature Human Immunodeficiency Virus Type 1 Particles. J. Virol. 74, 9381–9387 (2000).

35. Croft, J. T., et al. Reconstructing a Missing Link of HIV-1 Assembly: HIV-1 Envelope-Matrix Interactions in a Native Viral Context.

36. Brugger, B. et al. The HIV lipidome: a raft with an unusual composition. Proc Natl Acad Sci U A 103, 2641–6 (2006).

37. Lorizate, M. et al. Comparative lipidomics analysis of HIV-1 particles and their producer cell membrane in different cell lines. Cell. Microbiol. 15, 292–304 (2013).

38. Majumder, A. & Voth, G. A. Structural Heterogeneity of the Membrane-Interacting Region of the HIV-1 Envelope Glycoprotein. J. Am. Chem. Soc. 10.1021/jacs.5c15421 (2025) doi:10.1021/jacs.5c15421.

39. Shehata, M. et al. N-Glycans Modulate HIV-1 Env Conformational Plasticity. bioRxiv 2025.03.26.645577 (2025) doi:10.1101/2025.03.26.645577.

40. Souza, P. C. T. et al. Martini 3: a general purpose force field for coarse-grained molecular dynamics. Nat Methods 18, 382–388 (2021).

41. Ingólfsson, H. I. et al. The power of coarse graining in biomolecular simulations. Wiley Interdiscip. Rev. Comput. Mol. Sci. 4, 225–248 (2014).

42. Alessandri, R. et al. Pitfalls of the Martini Model. J. Chem. Theory Comput. 15, 5448–5460 (2019).

43. Marrink, S. J. et al. Computational Modeling of Realistic Cell Membranes. Chem. Rev. 119, 6184–6226 (2019).

44. Chen Steve S.-L., et al. Identification of the LWYIK Motif Located in the Human Immunodeficiency Virus Type 1 Transmembrane gp41 Protein as a Distinct Determinant for Viral Infection. J. Virol. 83, 870–883 (2009).

45. Mattei, S., Schur, F. K. & Briggs, J. A. Retrovirus maturation-an extraordinary structural transformation. Curr. Opin. Virol. 18, 27–35 (2016).

46. Pornillos, O. & Ganser-Pornillos, B. K. Maturation of retroviruses. Curr. Opin. Virol. 36, 47–55 (2019).

47. Qu, K. et al. Maturation of the matrix and viral membrane of HIV-1. Science 373, 700–704 (2021).

48. Murakami, T., Ablan, S., Freed, E. O. & Tanaka, Y. Regulation of Human Immunodeficiency Virus Type 1 Env-Mediated Membrane Fusion by Viral Protease Activity. J. Virol. 78, 1026–1031 (2004).

49. Jiang, J. & Aiken, C. Maturation-Dependent Human Immunodeficiency Virus Type 1 Particle Fusion Requires a Carboxyl-Terminal Region of the gp41 Cytoplasmic Tail. J. Virol. 81, 9999–10008 (2007).

50. Wyma, D. J. et al. Coupling of Human Immunodeficiency Virus Type 1 Fusion to Virion Maturation: a Novel Role of the gp41 Cytoplasmic Tail. J. Virol. 78, 3429–3435 (2004).

51. Chen, M. & Ludtke, S. J. Deep learning-based mixed-dimensional Gaussian mixture model for characterizing variability in cryo-EM. Nat. Methods 18, 930–936 (2021).

52. Lu, M. et al. Associating HIV-1 envelope glycoprotein structures with states on the virus observed by smFRET. Nature 568, 415–419 (2019).

53. Munro, J. B. et al. Conformational dynamics of single HIV-1 envelope trimers on the surface of native virions. Science 346, 759–763 (2014).

54. Ma, X. et al. HIV-1 Env trimer opens through an asymmetric intermediate in which individual protomers adopt distinct conformations. eLife 7, (2018).

55. Herschhorn, A. et al. Release of gp120 Restraints Leads to an Entry-Competent Intermediate State of the HIV-1 Envelope Glycoproteins. mBio 7, (2016).

56. Wang, Q., Finzi, A. & Sodroski, J. The Conformational States of the HIV-1 Envelope Glycoproteins. Trends Microbiol. 28, 655–667 (2020).

57. Cale, E. M. et al. Antigenic analysis of the HIV-1 envelope trimer implies small differences between structural states 1 and 2. J. Biol. Chem. 298, 101819 (2022).

58. Torralba, J. et al. Cholesterol Constrains the Antigenic Configuration of the Membrane-Proximal Neutralizing HIV-1 Epitope. ACS Infect. Dis. 6, 2155–2168 (2020).

59. Aloia, R. C., Tian, H. & Jensen, F. C. Lipid composition and fluidity of the human immunodeficiency virus envelope and host cell plasma membranes. Proc Natl Acad Sci U A 90, 5181–5 (1993).

60. Brügger, B. et al. The HIV lipidome: a raft with an unusual composition. Proc. Natl. Acad. Sci. U. S. A. 103, 2641–2646 (2006).

61. Greenwood, A. I. et al. CRAC motif peptide of the HIV-1 gp41 protein thins SOPC membranes and interacts with cholesterol. Biochim. Biophys. Acta 1778, 1120–1130 (2008).

62. Schwarzer, R. et al. The cholesterol-binding motif of the HIV-1 glycoprotein gp41 regulates lateral sorting and oligomerization. Cell. Microbiol. 16, 1565–1581 (2014).

63. Vishwanathan, S. A. et al. Hydrophobic substitutions in the first residue of the CRAC segment of the gp41 protein of HIV. Biochemistry 47, 124–130 (2008).

64. Leaman, D. P. & Zwick, M. B. Increased functional stability and homogeneity of viral envelope spikes through directed evolution. PLoS Pathog. 9, e1003184 (2013).

65. Souza, P. C. T. et al. GōMartini 3: From large conformational changes in proteins to environmental bias corrections. Nat. Commun. 16, 4051 (2025).

66. Leaman, D. P., Stano, A., Chen, Y., Zhang, L. & Zwick, M. B. Membrane Env Liposomes Facilitate Immunization with Multivalent Full-Length HIV Spikes. J. Virol. 95, e0000521 (2021).

67. Mastronarde, D. N. SerialEM: A Program for Automated Tilt Series Acquisition on Tecnai Microscopes Using Prediction of Specimen Position. Microsc. Microanal. 9, 1182–1183 (2003).

68. Zheng, S. Q. et al. MotionCor2: anisotropic correction of beam-induced motion for improved cryo-electron microscopy. Nat. Methods 14, 331–332 (2017).

69. Kremer, J. R., Mastronarde, D. N. & McIntosh, J. R. Computer Visualization of Three-Dimensional Image Data Using IMOD. J. Struct. Biol. 116, 71–76 (1996).

70. Chen, M. et al. A complete data processing workflow for cryo-ET and subtomogram averaging. Nat. Methods 16, 1161–1168 (2019).

71. Bepler, T., Kelley, K., Noble, A. J. & Berger, B. Topaz-Denoise: general deep denoising models for cryoEM and cryoET. Nat. Commun. 11, 5208 (2020).

72. Pettersen, E. F. et al. UCSF Chimera--a visualization system for exploratory research and analysis. J. Comput. Chem. 25, 1605–1612 (2004).

73. Meng, E. C. et al. UCSF ChimeraX: Tools for structure building and analysis. Protein Sci. Publ. Protein Soc. 32, e4792 (2023).

74. Kluyver, T. et al. Jupyter Notebooks–a publishing format for reproducible computational workflows. in Positioning and power in academic publishing: Players, agents and agendas 87–90 (IOS press, 2016).

75. Waterhouse, A. et al. SWISS-MODEL: homology modelling of protein structures and complexes. Nucleic Acids Res. 46, W296–W303 (2018).

76. Abraham, M. J. et al. GROMACS: High performance molecular simulations through multi-level parallelism from laptops to supercomputers. SoftwareX 1–2, 19–25 (2015).

77. Case, D. A. Amber 2025. (2025).

78. Gowers, R. J. et al. MDAnalysis: A Python Package for the Rapid Analysis of Molecular Dynamics Simulations. in (Los Alamos National Laboratory (LANL), Los Alamos, NM (United States), 2019). doi:10.25080/Majora-629e541a-00e.

79. Michaud-Agrawal, N., Denning, E. J., Woolf, T. B. & Beckstein, O. MDAnalysis: a toolkit for the analysis of molecular dynamics simulations. J. Comput. Chem. 32, 2319–2327 (2011).

80. Park, S.-J. et al. CHARMM-GUI Glycan Modeler for modeling and simulation of carbohydrates and glycoconjugates. Glycobiology 29, 320–331 (2019).

81. Wu, E. L. et al. CHARMM-GUI Membrane Builder toward realistic biological membrane simulations. J. Comput. Chem. 35, 1997–2004 (2014).

82. Jo, S., Kim, T., Iyer, V. G. & Im, W. CHARMM-GUI: a web-based graphical user interface for CHARMM. J. Comput. Chem. 29, 1859–1865 (2008).

83. Cao, L. et al. Global site-specific N-glycosylation analysis of HIV envelope glycoprotein. Nat. Commun. 8, 14954 (2017).

84. Huang, J. et al. CHARMM36m: an improved force field for folded and intrinsically disordered proteins. Nat. Methods 14, 71–73 (2017).

85. Hess, B., Bekker, H., Berendsen, H. J. C. & Fraaije, J. G. E. M. LINCS: A linear constraint solver for molecular simulations. J. Comput. Chem. 18, 1463–1472 (1997).

86. Bussi, G., Donadio, D. & Parrinello, M. Canonical sampling through velocity rescaling. J. Chem. Phys. 126, 014101 (2007).

87. Bernetti, M. & Bussi, G. Pressure control using stochastic cell rescaling. J. Chem. Phys. 153, 114107 (2020).

88. Do, H. N. & Gnanakaran, S. Iterative Multiscale Molecular Dynamics: Accelerating Conformational Sampling of Biomolecular Systems by Iterating All-Atom and Coarse-Grained Molecular Dynamics Simulations. bioRxiv 10.1101/2025.02.16.638568 (2025) doi:10.1101/2025.02.16.638568.

89. Kroon, P. C. et al. Martinize2 and Vermouth: Unified Framework for Topology Generation. 10.7554/elife.90627.3 (2025) doi:10.7554/elife.90627.3.

90. Koukos, P. I. et al. Martini 3 Force Field Parameters for Protein Lipidation Post-Translational Modifications. J. Chem. Theory Comput. 19, 8901–8918 (2023).

91. Wassenaar, T. A., Pluhackova, K., Böckmann, R. A., Marrink, S. J. & Tieleman, D. P. Going Backward: A Flexible Geometric Approach to Reverse Transformation from Coarse Grained to Atomistic Models. J. Chem. Theory Comput. 10, 676–690 (2014).

92. Herzog, F. A., Braun, L., Schoen, I. & Vogel, V. Improved Side Chain Dynamics in MARTINI Simulations of Protein–Lipid Interfaces. J. Chem. Theory Comput. 12, 2446–2458 (2016).

93. Periole, X., Cavalli, M., Marrink, S.-J. & Ceruso, M. A. Combining an Elastic Network With a Coarse-Grained Molecular Force Field: Structure, Dynamics, and Intermolecular Recognition. J. Chem. Theory Comput. 5, 2531–2543 (2009).

94. Wassenaar, T. A., Ingólfsson, H. I., Böckmann, R. A., Tieleman, D. P. & Marrink, S. J. Computational Lipidomics with insane: A Versatile Tool for Generating Custom Membranes for Molecular Simulations. J. Chem. Theory Comput. 11, 2144–2155 (2015).

95. Tironi, I. G., Sperb, R., Smith, P. E. & van Gunsteren, W. F. A generalized reaction field method for molecular dynamics simulations. J. Chem. Phys. 102, 5451–5459 (1995).

96. Grubmüller, H., Heller, H., Windemuth, A. & Schulten, K. Generalized Verlet Algorithm for Efficient Molecular Dynamics Simulations with Long-range Interactions. Mol. Simul. 6, 121–142 (1991).

97. Galaz-Montoya, J. G., Flanagan, J., Schmid, M. F. & Ludtke, S. J. Single particle tomography in EMAN2. J. Struct. Biol. 190, 279–290 (2015).

98. Croll, T. I. ISOLDE: a physically realistic environment for model building into low-resolution electron-density maps. Acta Crystallogr. Sect. Struct. Biol. 74, 519–530 (2018).

99. Zhang, Z., Zhang, A. & Xiao, G. Improved Protein Hydrogen/Deuterium Exchange Mass Spectrometry Platform with Fully Automated Data Processing. Anal. Chem. 84, 4942–4949 (2012).

100. Watson, M. J. et al. Simple Platform for Automating Decoupled LC–MS Analysis of Hydrogen/Deuterium Exchange Samples. J. Am. Soc. Mass Spectrom. 32, 597–600 (2021).

101. Fang, J., Rand, K. D., Beuning, P. J. & Engen, J. R. False EX1 signatures caused by sample carryover during HX MS analyses. Int. J. Mass Spectrom. 302, 19–25 (2011).

102. Li, Z. et al. Subnanometer structures of HIV-1 envelope trimers on aldrithiol-2-inactivated virus particles. Nat. Struct. Amp Mol. Biol. 27, 726—734 (2020).

