## Supplementary Figs 1-11 for "Structure of HIV-1 Env glycoprotein on virions reveals an alternative fusion subunit organization and native membrane coupling"

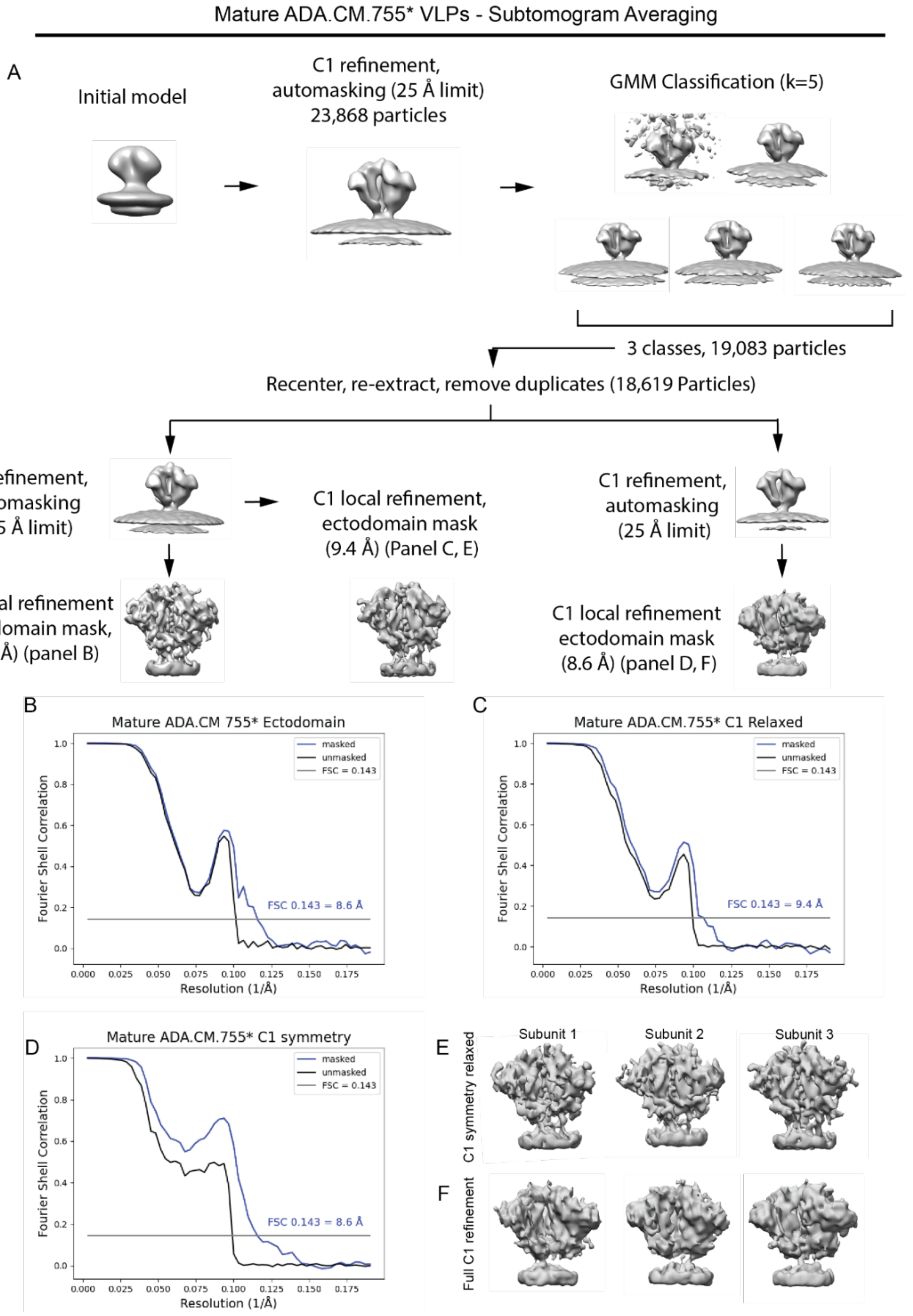


**Figure S1. Subtomogram averaging of mature ADA.CM.755* ectodomain. (A)** Subtomogram averaging pipeline for the mature ADA.CM.755* Env ectodomain, as described in the methods.

**(B)** Fourier Shell Correlation (FSC) for the mature ADA.CM.755* Env focused refinement of the ectodomain with C3 symmetry. Using FSC = 0.143 criterion, we report 8.6 Å resolution. **(C)** FSC for the mature ADA.CM.755* Env focused refinement of the ectodomain with symmetry relaxation to C1. Using FSC = 0.143 criterion, we report 9.4 Å resolution. **(D)** FSC for the mature ADA.CM.755* Env focused refinement of the ectodomain with C1 symmetry. Using FSC = 0.143 criterion, we report 8.6 Å resolution. Throughout the figure, the blue curves represent the masked FSC and the black curves represent the FSC calculated with unmasked maps. **(E, F)** Comparison of different subunits between the C1 relaxed **(E)** and Full C1 refinements **(F)**.

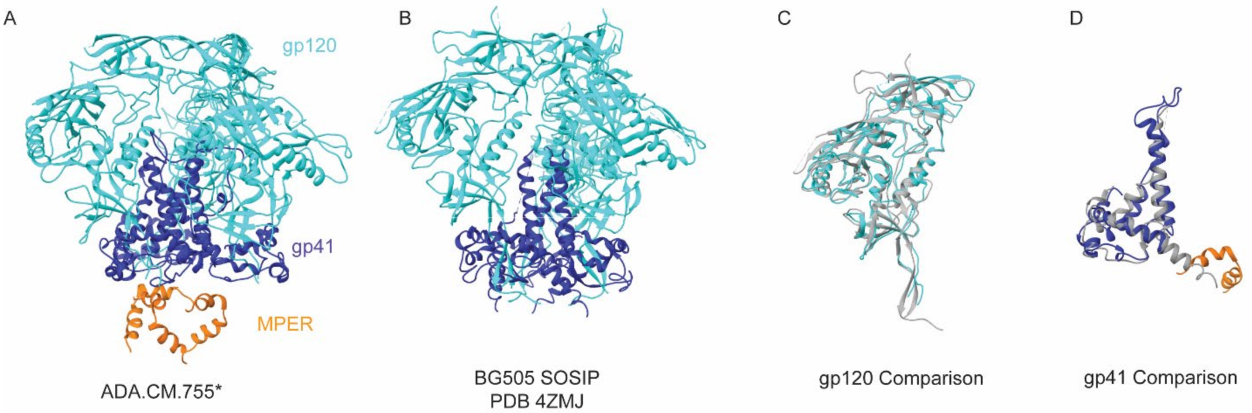


**Figure S2. Comparison of ADA.CM.755* Env structure determined by subtomogram averaging to BG505.SOSIP. (A)** Homology model of ADA.CM.755* Env was created using SWISS-MODEL and fitted to the cryo-ET density map as described in the methods. Throughout the figure, gp120 is colored cyan, gp41 ectodomain is colored blue, and the MPER region of gp41 is colored orange. **(B)** Model of BG505.SOSIP Env (PDB: 4ZMJ) [1]. **(C)** Comparison of gp120 in ADA.CM.755* Env (cyan) to BG505 (gray) **(D)** Comparison of gp41 in ADA.CM.755* Env (blue, orange) to BG505 (gray). The α-9 helix is not modeled in the ADA.CM.755* Env.

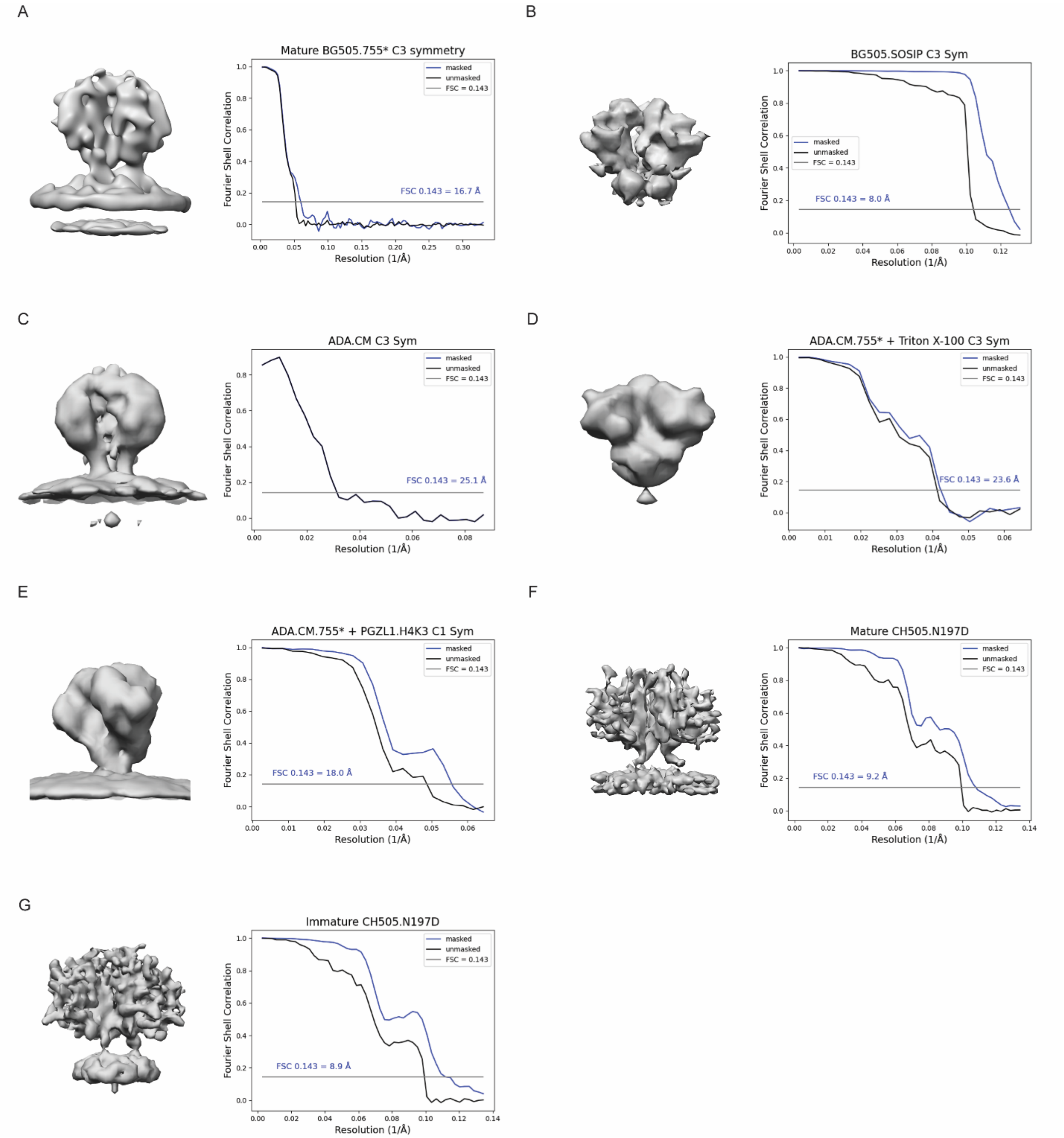


**Figure S3. Fourier Shell Correlations (FSC) for a panel of Env reconstructions by subtomogram averaging. (A)** BG505.755* Env on the viral membrane with C3 symmetry applied. Using FSC = 0.143 criterion, we report 16.7 Å resolution. **(B)** BG505.SOSIP with C3 symmetry applied. Using FSC = 0.143 criterion, we report 8.0 Å resolution. **(C)** ADA.CM Env with C3 symmetry applied. Using FSC = 0.143 criterion, we report 25.1 Å resolution. **(D)** ADA.CM.755* Env on membrane-stripped PR55$^{Gag}$ lattices treated with Triton X-100. C3 symmetry applied. Using FSC = 0.143 criterion, we report 23.6 Å resolution. **(E)** ADA.CM.755*

Env treated with PGLZ1.H4K3 Fab. C1 Symmetry applied. Using FSC = 0.143 criterion, we report 18.0 Å resolution. **(F)** Mature CH505.N197D Env. C3 Symmetry applied. Using FSC = 0.143 criterion, we report 9.2 Å resolution. **(G)** Immature CH505.N197D Env. C3 Symmetry applied. Using FSC = 0.143 criterion, we report 8.9 Å resolution.

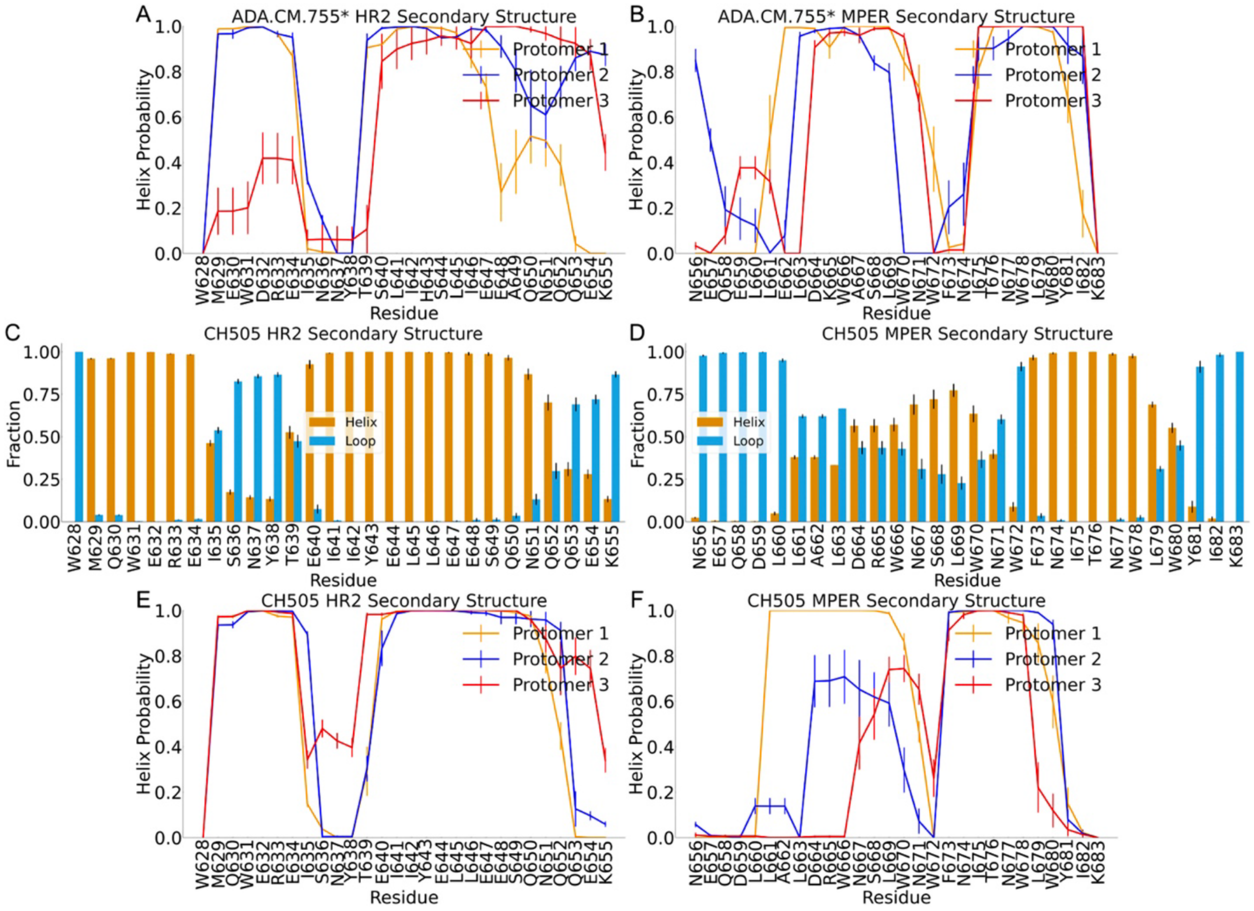


**Figure S4. Secondary structures of the HR2 and MPER residues sampled from the MD simulations of HIV-1 Env. (A-B)** Differences in the helix probability of HR2 **(A)** and MPER **(B)** residues from different protomers of ADA.CM.755* sampled from the MD simulations of ADA.CM.755*. **(C-D)** Secondary structures of the **(C)** HR2 and **(D)** MPER residues sampled from the MD simulations (both density-guided and unbiased) of HIV-1 Env CH505. The fractions of simulation time the residues existed as helices are colored blue, while the fractions of simulation time the residues existed as loops are colored orange. **(E-F)** Differences in the helix probability of HR2 **(A)** and MPER **(B)** residues from different protomers of CH505 sampled from the MD simulations of CH505. The error bars represent the standard errors of the means (S.E.m.) from different simulation replicas.

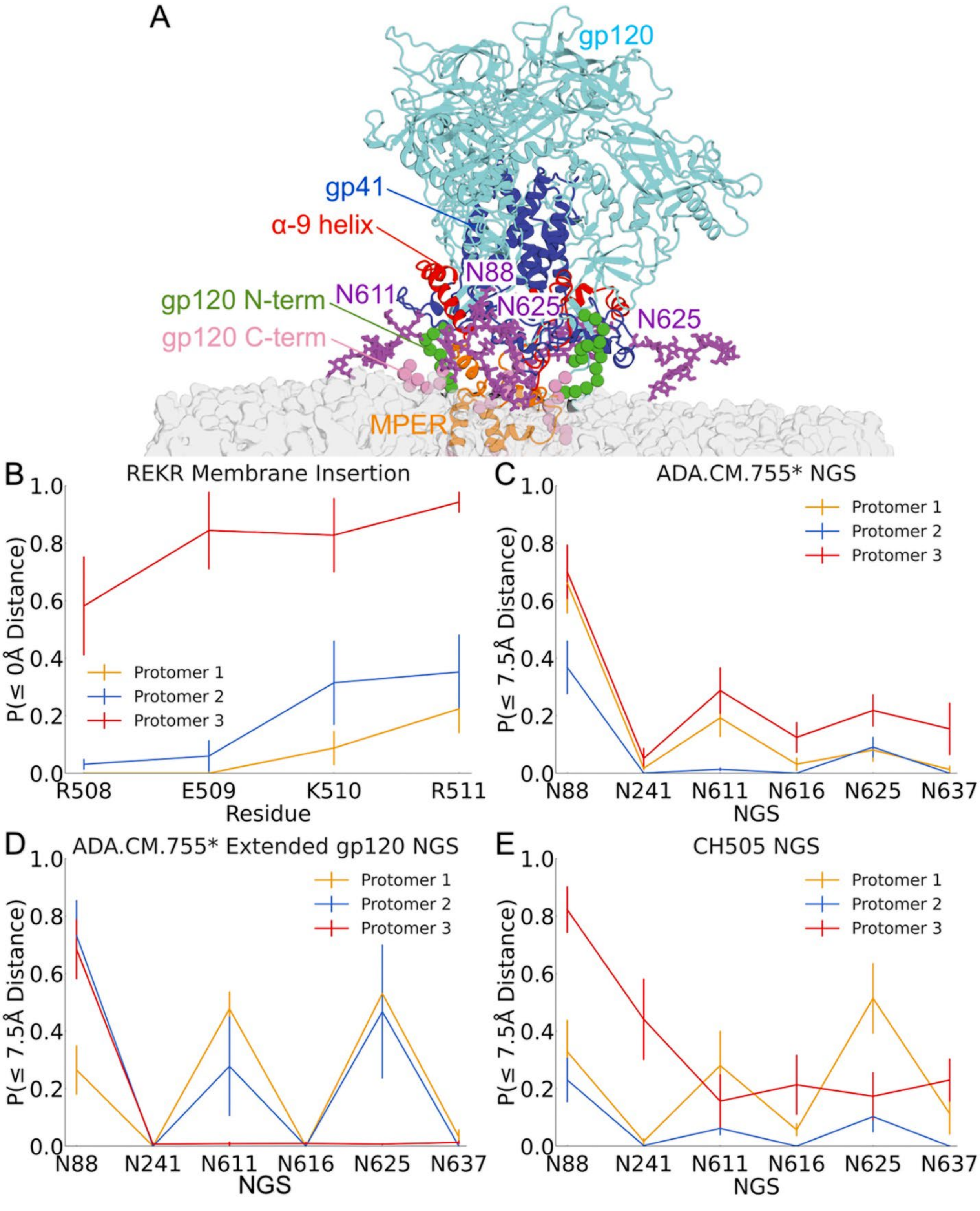


**Figure S5. Interactions of the gp120 C-terminal residues (REKR) and N-glycosylation sites (NGS) with the membrane captured from the MD simulations. (A)** Representative conformation of the insertion of the extended gp120 C-termini of ADA.CM.755* and glycan interactions with the membrane captured from the MD simulations. In this conformation, one glycan interaction from residue N88 and two glycan interactions from both residues N611 (one eclipsed on the other side of the image) and N625 were also observed. The gp120 domain is colored cyan, gp41 ectodomain is colored blue, extended gp120 N-terminus is colored green, gp41 HR2 is colored red, MPER is colored orange, most glycans are hidden, except those interacting with membrane colored purple, and the membrane is colored gray. **(B)** Fractions of simulation times the extended gp120 C-terminus residues R508, E509, K510, and R511 located on the same or below the plane (≤ 0Å distance) formed by the phosphate lipid headgroups in the upper leaflet. **(C-E)** Fractions of simulation times the glycans attached to NGS residues N88, N241, N611, N616, N625, and N637 located at or below 7.5 Å (≤ 7.5Å distance) to the plane formed by the phosphate lipid headgroups in the upper leaflet sampled from the MD simulations of ADA.CM.755* **(C)**, ADA.CM.755* with extended gp120 termini **(D)**, and glycosylated CH505 **(E)**. Here, only NGS with average glycan heights of ≤ 20 Å in at least one protomer were included. The error bars represent the S.E.m. from different simulation replicas.

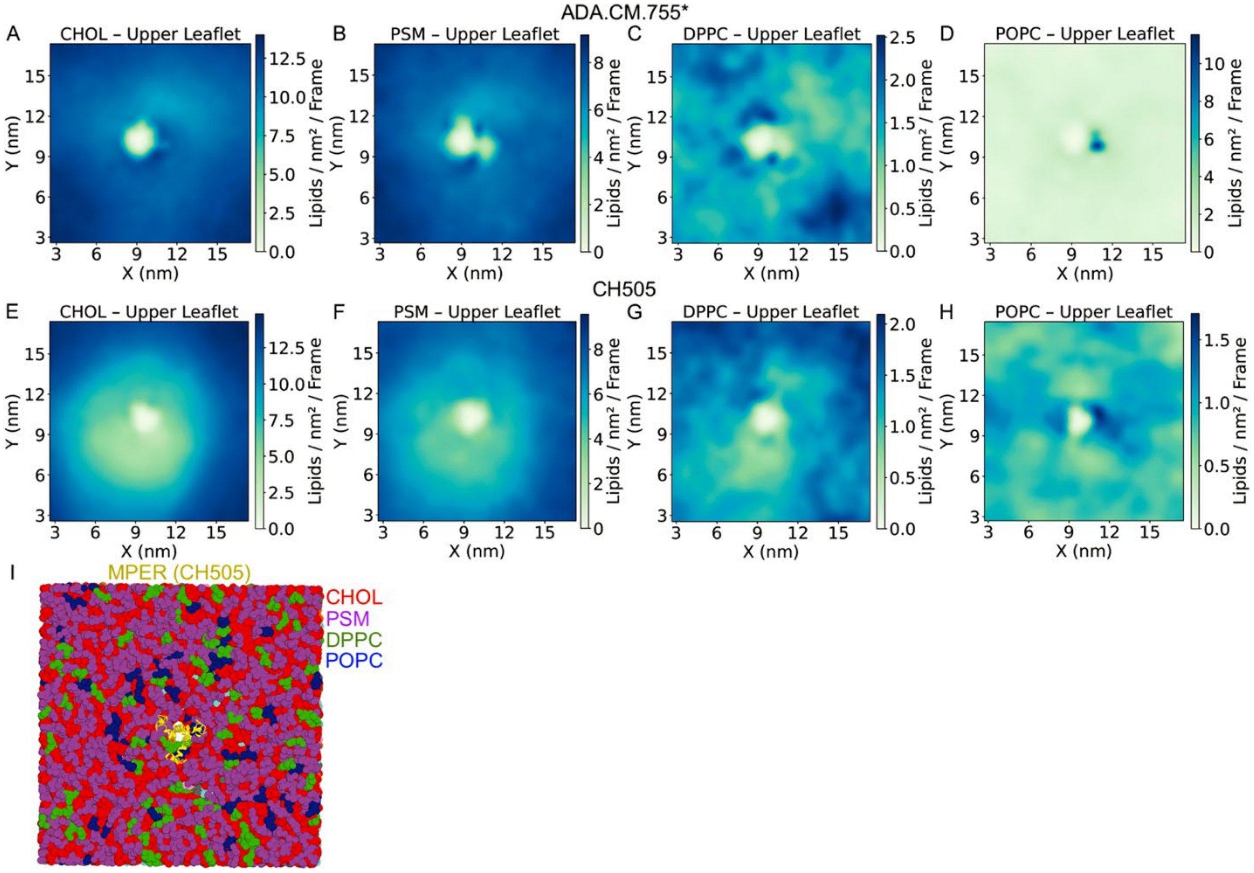


**Figure S6. Distributions and representative conformations of lipids in the upper leaflet during the CG MD simulations. (A-H)** Distributions of the four types of lipid molecules, including CHOL, PSM, DPPC, and POPC, in the upper leaflet of the HIV-1 membrane observed in the last 5μs of CG simulations of ADA.CM.755* with extended gp120 termini and MA **(A-D)** and mature CH505 **(E-H)**. A color scheme of green – blue was used to illustrate the lowest – highest average density of the lipid molecules in the lower leaflet in the simulations. **(I)** Representative conformation of CH505 in the HIV-1 membrane. Since the HIV-1 proteins were completely restrained during the CG simulations, the atomistic conformations of the MPER before the CG simulations were used for better visualizations. The HIV-1 proteins are colored yellow, CHOL lipid molecules are colored red, DPPC are colored green, POPC are colored blue, and PSM are colored magenta.

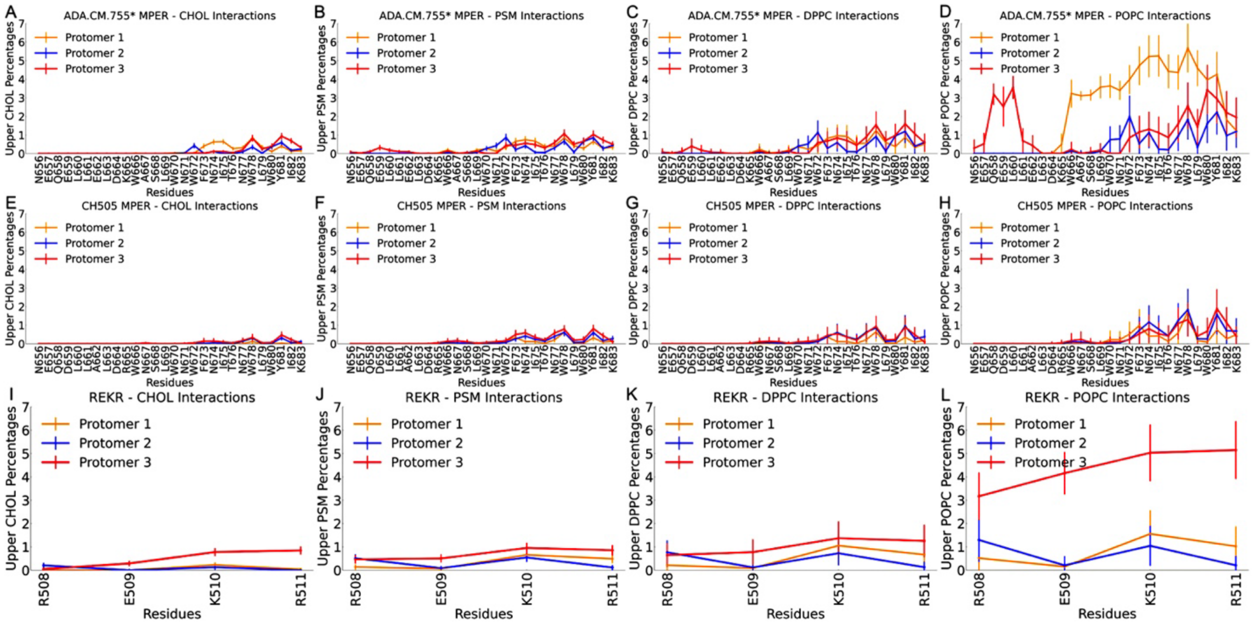


**Figure S7. Interactions of the MPER and gp120 termini (R508-R511) residues with upper leaflet lipid molecules determined from the last 5μs of CG simulations of HIV-1 Env in the HIV-1 membrane. (A-D)** Interactions of MPER residues from different protomers of ADA.CM.755* with CHOL **(A)**, PSM **(B)**, DPPC **(C)**, and POPC **(D)** sampled from the CG simulations of the HIV-1 Env in the HIV-1 membrane. **(E-H)** Interactions of MPER residues from different protomers of CH505 with CHOL **(E)**, PSM **(F)**, DPPC **(G)**, and POPC **(H)** sampled from the CG simulations of the HIV-1 Env in the HIV-1 membrane. **(I-L)** Interactions of gp120 termini (R508-R511) residues from different protomers with CHOL **(I)**, PSM **(J)**, DPPC **(K)**, and POPC **(L)** sampled from the CG simulations of ADA.CM.755* in the HIV-1 membrane. The error bars represent the S.E.m. from different simulation replicas.

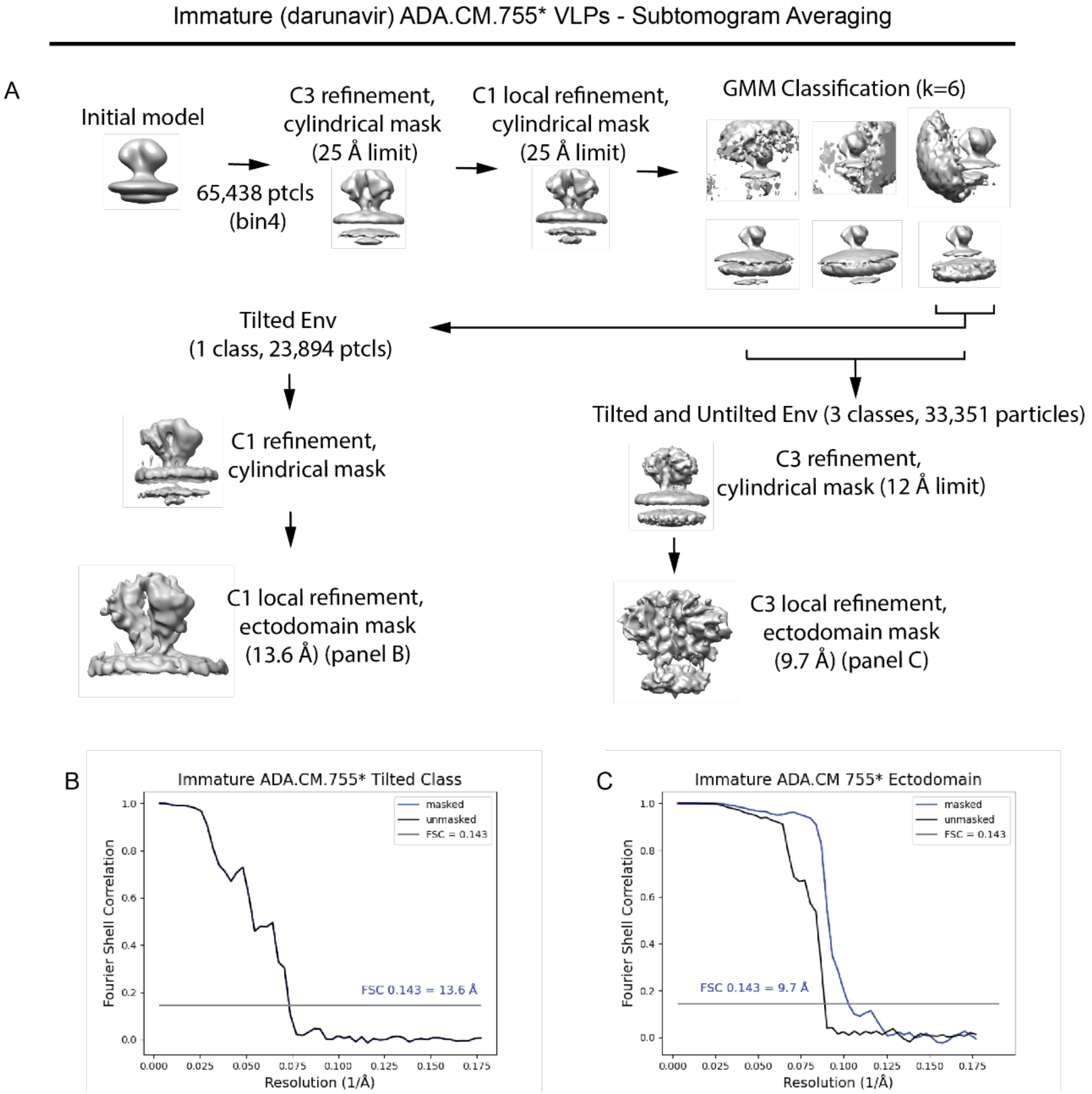


**Figure S8. Subtomogram averaging of immature ADA.CM.755* ectodomain. (A)** Subtomogram averaging pipeline for the immature ADA.CM.755* Env ectodomain, as described in the methods. **(B)** Fourier Shell Correlation (FSC) for the tilted immature ADA.CM.755* Env focused refinement of the ectodomain with C1 symmetry. Using FSC = 0.143 criterion, we report 13.6 Å resolution. **(C)** Fourier Shell Correlation (FSC) for the immature ADA.CM.755* Env focused refinement of the ectodomain with C3 symmetry. Using FSC = 0.143 criterion, we report 9.6 Å resolution.

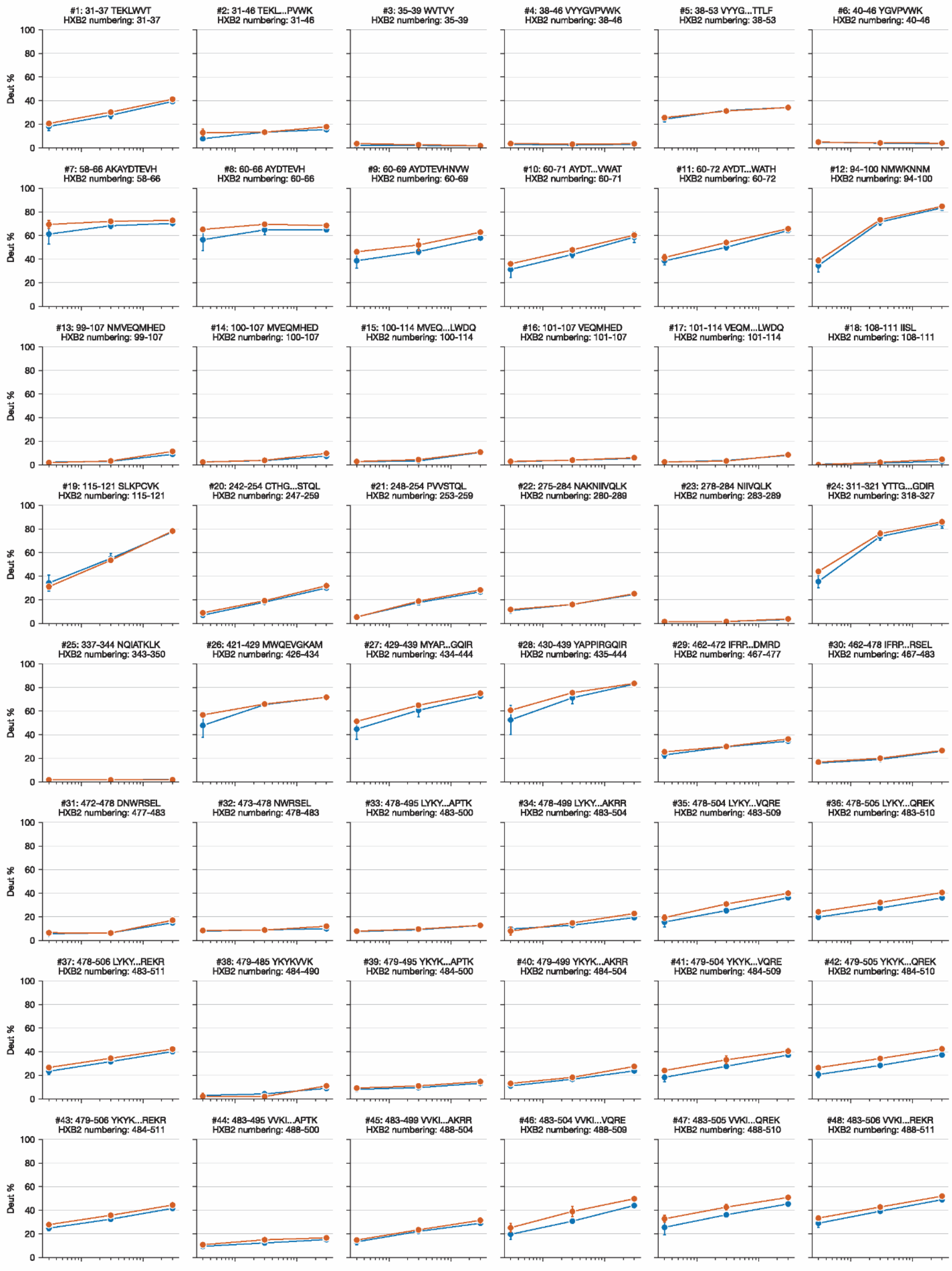
Deut %
100
80
60
40
20
0
#1: 31-37 TEKLWVT
HXB2 numbering: 31-37
#2: 31-46 TEKL...PVWK
HXB2 numbering: 31-46
#3: 35-39 WVTVY
HXB2 numbering: 35-39
#4: 38-46 VYYGVPVWK
HXB2 numbering: 38-46
#5: 38-53 VYYG...TTLF
HXB2 numbering: 38-53
#6: 40-46 YGVPVWK
HXB2 numbering: 40-46
#7: 58-66 AKAYDTEVH
HXB2 numbering: 58-66
#8: 60-66 AYDTEVH
HXB2 numbering: 60-66
#9: 60-69 AYDTEVHNVW
HXB2 numbering: 60-69
#10: 60-71 AYDT...VWAT
HXB2 numbering: 60-71
#11: 60-72 AYDT...WATH
HXB2 numbering: 60-72
#12: 94-100 NMWKNNM
HXB2 numbering: 94-100
#13: 99-107 NMVEQMHED
HXB2 numbering: 99-107
#14: 100-107 MVEQMHED
HXB2 numbering: 100-107
#15: 100-114 MVEQ...LWDQ
HXB2 numbering: 100-114
#16: 101-107 VEQMHED
HXB2 numbering: 101-107
#17: 101-114 VEQM...LWDQ
HXB2 numbering: 101-114
#18: 108-111 IISL
HXB2 numbering: 108-111
#19: 115-121 SLKPCVK
HXB2 numbering: 115-121
#20: 242-254 CTHG...STQL
HXB2 numbering: 247-259
#21: 248-254 PVVSTQL
HXB2 numbering: 253-259
#22: 275-284 NAKNIIVQLK
HXB2 numbering: 280-289
#23: 278-284 NIIVQLK
HXB2 numbering: 283-289
#24: 311-321 YTTG...GDIR
HXB2 numbering: 318-327
#25: 337-344 NQIATKLK
HXB2 numbering: 343-350
#26: 421-429 MWQEVGKAM
HXB2 numbering: 426-434
#27: 429-439 MYAP...GQIR
HXB2 numbering: 434-444
#28: 430-439 YAPPIRGQIR
HXB2 numbering: 435-444
#29: 462-472 IFRP...DMRD
HXB2 numbering: 467-477
#30: 462-478 IFRP...RSEL
HXB2 numbering: 467-483
#31: 472-478 DNWRSEL
HXB2 numbering: 477-483
#32: 473-478 NWRSEL
HXB2 numbering: 478-483
#33: 478-495 LYKY...APTK
HXB2 numbering: 483-500
#34: 478-499 LYKY...AKRR
HXB2 numbering: 483-504
#35: 478-504 LYKY...VQRE
HXB2 numbering: 483-509
#36: 478-505 LYKY...QREK
HXB2 numbering: 483-510
#37: 478-506 LYKY...REKR
HXB2 numbering: 483-511
#38: 479-485 YKYKVVK
HXB2 numbering: 484-490
#39: 479-495 YKYK...APTK
HXB2 numbering: 484-500
#40: 479-499 YKYK...AKRR
HXB2 numbering: 484-504
#41: 479-504 YKYK...VQRE
HXB2 numbering: 484-509
#42: 479-505 YKYK...QREK
HXB2 numbering: 484-510
#43: 479-506 YKYK...REKR
HXB2 numbering: 484-511
#44: 483-495 VVKI...APTK
HXB2 numbering: 488-500
#45: 483-499 VVKI...AKRR
HXB2 numbering: 488-504
#46: 483-504 VVKI...VQRE
HXB2 numbering: 488-509
#47: 483-505 VVKI...QREK
HXB2 numbering: 488-510
#48: 483-506 VVKI...REKR
HXB2 numbering: 488-511

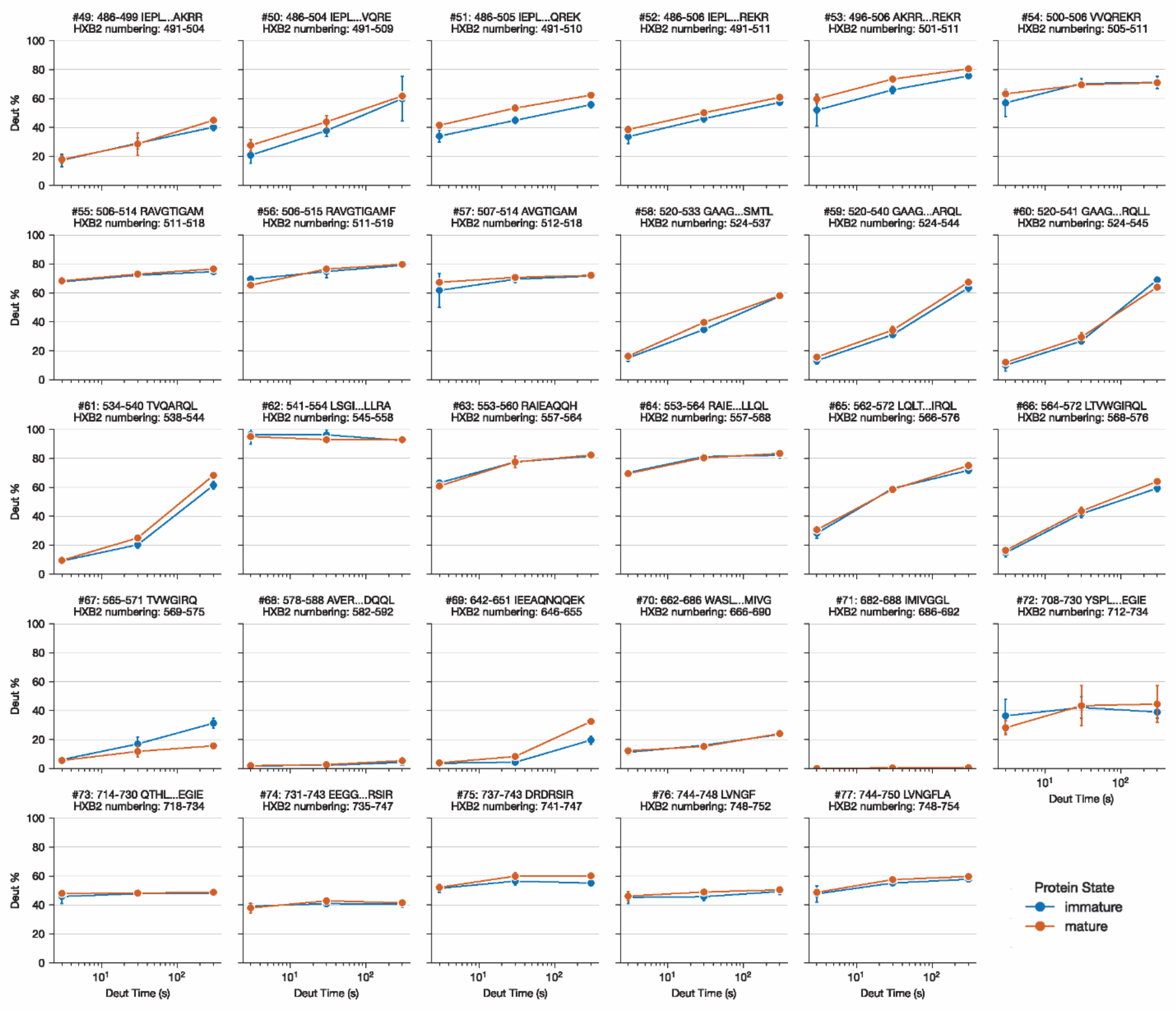


**Figure S9. Deuterium uptake plots from HDX-MS of HIV-1 Env.** Deuterium uptake plots for all peptides in Env ectodomain commonly identified between ADA.CM.755* Env from mature VLPs (orange) and immature VLPs (blue). The sequence of the peptides are given above each uptake plot using HXB2 numbering.

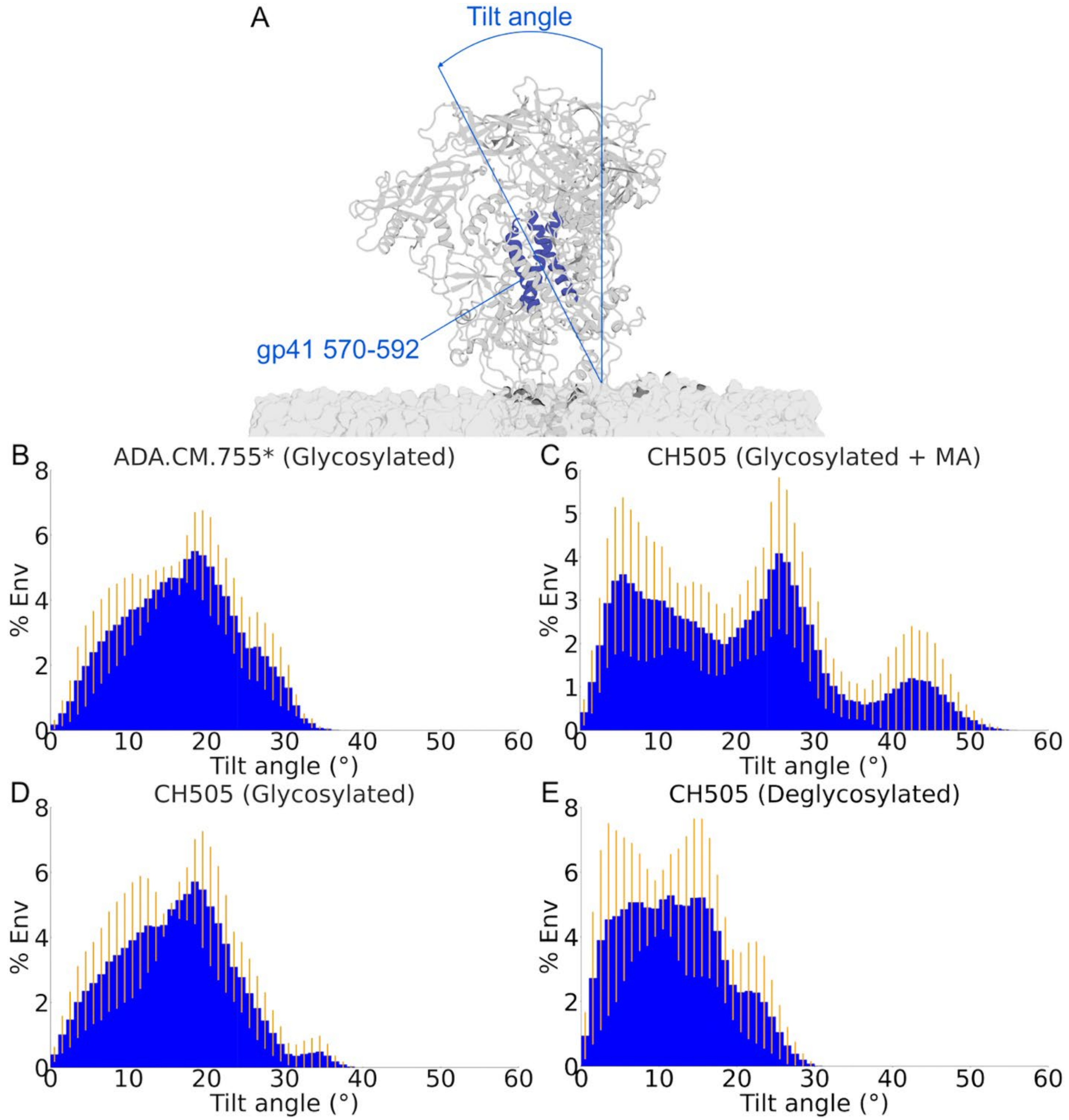


**Figure S10. Distributions of the tilt angles of the ectodomains sampled from the MD simulations of HIV-1 Env. (A)** The tilt angles of the ectodomains were calculated by measuring the angles formed by the vector drawn through the center-of-mass (COM) of the gp41 residues 570-592 (blue), the outer membrane center, and the z-axis. **(B-E)** Histograms of tilt angles of the glycosylated ADA.CM.755* without MA **(B)** and the CH505 envelope glycoproteins in its glycosylated form with MA **(C)**, in its glycosylated form without MA **(D)**, and in its deglycosylated form **(E)**. The error bars represent the S.E.m. from different simulation replicas.

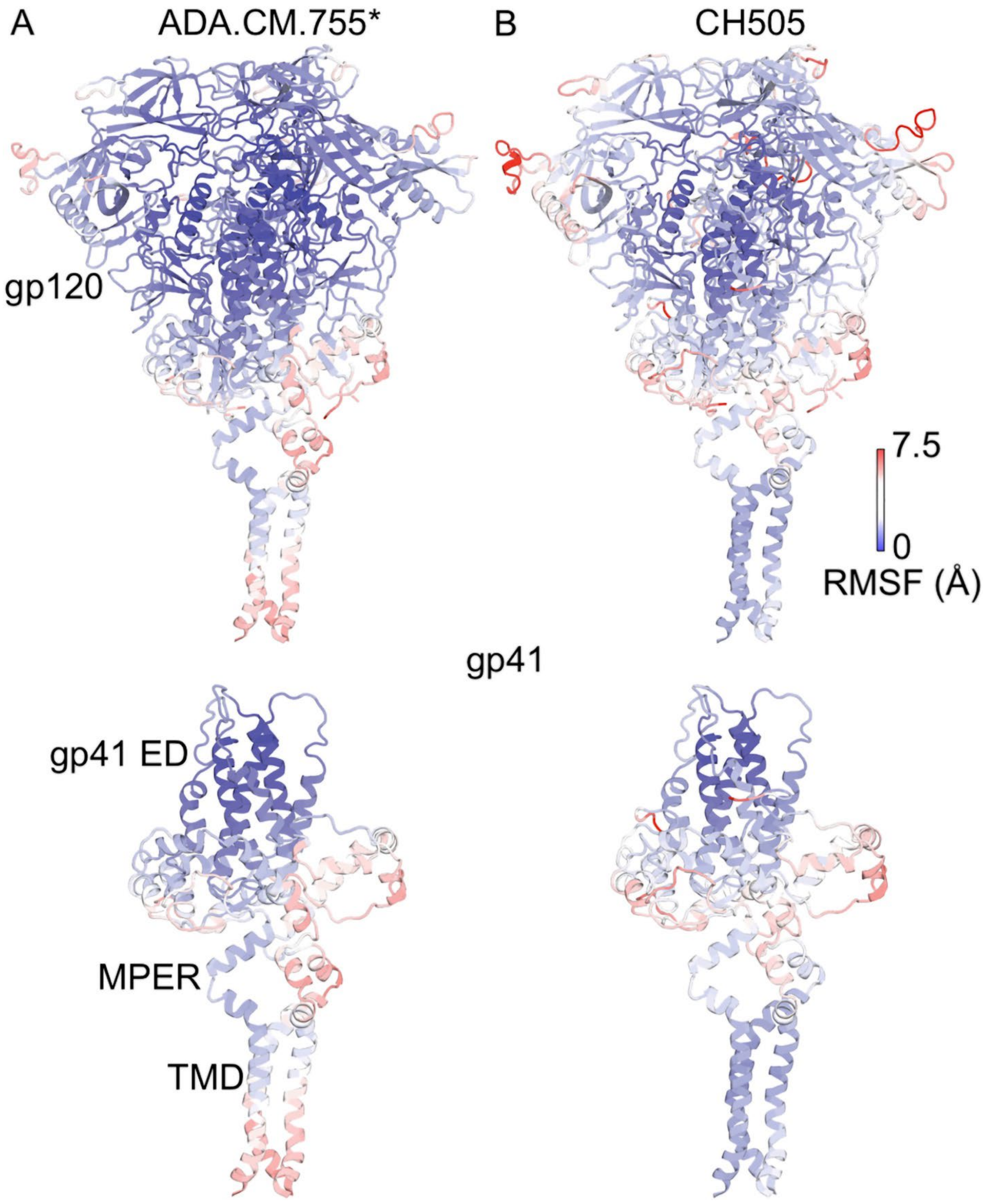


**Figure S11. Flexibility of the HIV-1 Env glycoproteins sampled from the MD simulations of ADA.CM.755* and glycosylated CH505.** The Env were shown up until the end of the gp41 TMD (HXB2 residue 708). The root-mean-square fluctuations (RMSF) of the backbone atoms were used to illustrate the flexibility of the HIV-1 Env. In particular, the Env RMSF was calculated for each simulation replica for each system, then the average Env RMSF across simulation replicas was calculated and shown in the figure. A color scale of blue (0 Å) – white – red (7.5 Å) was used to show the RMSF of both Env.

### References


1. Kwon, Y. D. *et al.* Crystal structure, conformational fixation and entry-related interactions of mature ligand-free HIV-1 Env. *Nat Struct Mol Biol* **22**, 522–531 (2015).
